# MALDI mass spectrometry imaging of fresh and processed food: constituents, ingredients, contaminants and additives

**DOI:** 10.1101/2021.12.23.473956

**Authors:** Julia Kokesch-Himmelreich, Oliver Wittek, Alan M. Race, Sophie Rakete, Claus Schlicht, Ulrich Busch, Andreas Römpp

## Abstract

Mass Spectrometry imaging (MS imaging) provides spatial information for a wide range of compound classes in different sample matrices. We used MS imaging to investigate the distribution of components in fresh and processed food, including meat, dairy and bakery products. The MS imaging workflow was optimized to cater to the specific properties and challenges of the individual samples. We successfully detected highly nonpolar and polar constituents such as beta-carotene and anthocyanins, respectively. For the first time, the distribution of a contaminant and a food additive was visualized in processed food. We detected acrylamide in German gingerbread and investigated the penetration of the preservative natamycin into cheese. For this purpose, a new data analysis tool was developed to study the penetration of analytes from uneven surfaces. Our results show that MS imaging has great potential in food analysis to provide relevant information about components’ distributions, particularly those underlying official regulations.

**Highlights:**

- Investigation of fresh and processed food by MALDI mass spectrometry imaging
- Visualization of different compound classes in plant and meat-based food
- Development of data processing tool for penetration/diffusion analysis (in food)
- Natamycin penetration in cheese, first visualization of food additive by MS imaging
- Acrylamide in gingerbread, first visualization of contaminant by MS imaging

## 1. Introduction

Mass spectrometry is extensively used for the analysis of food (Domínguez, Garrido Frenich, & Romero-González, 2020; Medina, Pereira, Silva, Perestrelo, & Câmara, 2019). It is ideally suited to address the complexity of food samples and to quantify a wide range of compounds. These studies are almost exclusively based on homogenized samples, which are commonly separated by chromatography before mass spectral detection.

However, in some cases, the identification and quantification of compounds is not sufficient, and the spatial distribution of one or more analytes is also relevant. The combination of mass spectrometry and spatial information is accessible by mass spectrometry imaging (MS imaging), which has gained substantial interest over the last 20 years in the analytical community, but has so far not been fully explored for food analysis. MS imaging enables the visualization of spatial distributions for a wide range of chemical compounds in complex biological samples (Römpp & Spengler, 2013; Spengler, 2015). The most widely used ionization technique is matrix-assisted laser desorption/ionization (MALDI).

The MALDI MS imaging measurement procedure is briefly explained by the example of a hardy kiwi (*Actinidia arguta*) in Figure 1. Commonly, thin sections of the sample are obtained and sprayed with a suitable matrix solution to obtain the incorporation of the analytes in small matrix crystals (co-crystallization). The matrix-coated surface of the sample is scanned in a grid-like pattern with a laser to desorb and ionize the analytes (Figure 1A). For each laser spot on the sample section, a full mass spectrum is acquired (segments shown in Figure 1B). An ‘ion image’ or ‘MS image’ can be generated by picking an *m/z* value of interest and displaying all pixels containing the selected signal in a predefined color, where the brightness of the color corresponds to the intensity of the ion signal (Figure 1C). Thus, the combination of mass spectrometric detection and spatially resolved analysis provides information about both the presence and the relative intensity of an analyte on the sample surface. Commonly, up to three ion channels are overlaid yielding a multicolor ion image (RGB MS image, Figure 1D, right), which can be directly compared to the optical image (Figure 1D, left). In the present example, the signals of a disaccharide, a phytochemical and a triglyceride, all compounds of interest in food analysis, were chosen to be visualized in the kiwi fruit. Details on these food constituents and their identification are discussed in the Results section. This example shows that the combination of MS images and the corresponding optical image can be used to discuss the potential function or effect of a given analyte. It is important to note that different compound classes can be visualized simultaneously - and without the need for labelling - despite their potentially varying physical and chemical properties.

**Figure 1:**
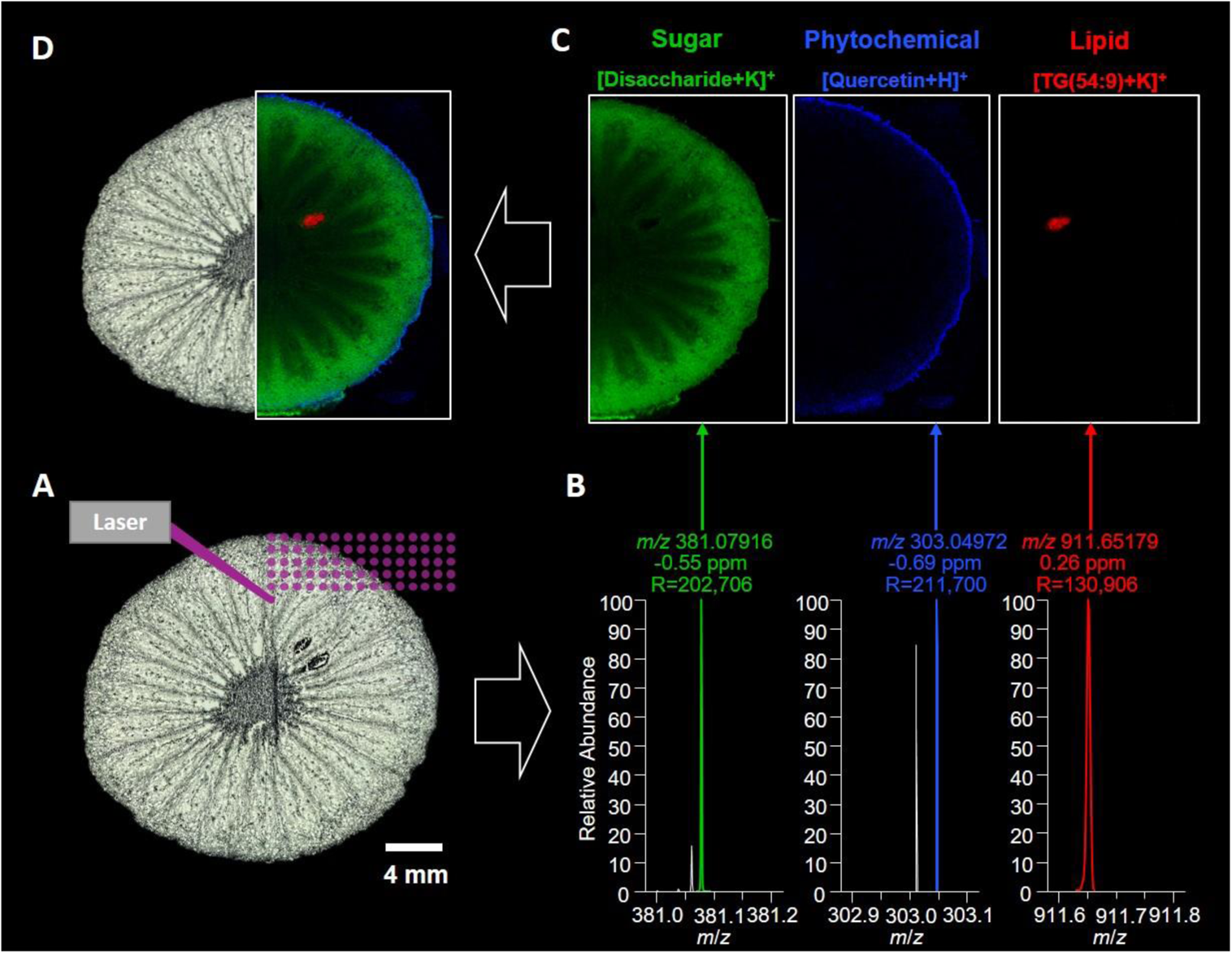
MS imaging of hardy kiwi. A: Optical image of kiwi section and schematic explanation of the MS imaging process, B: Single pixel mass spectra of three chosen analytes with their corresponding mass deviation in ppm and mass resolution R (FWHM). C: Single ion images of three chosen analytes, pixel size 45 µm, MS images were normalized to the total ion current (TIC). D: RGB MS image of the lipid TG(54:9 ([M+K]^+^, m/z 911.65255, red), Disaccharide ([M+K]^+^, m/z 381.07937, green), Quercetin ([M+H]^+^, m/z 303.04993, green).

MALDI MS imaging in general has been successfully used for analyzing metabolites (Sturtevant, Lee, & Chapman, 2016), drug compounds (Schulz, Becker, Groseclose, Schadt, & Hopf, 2019), lipids (Trim, Atkinson, Princivalle, Marshall, West, & Clench, 2008) and proteins (Meding et al., 2012) in biological samples. MS imaging methods were also developed for analyzing constituents in fresh plant-based food, e.g. wheat grain (Bhandari, Wang, Friedt, Spengler, Gottwald, & Römpp, 2015) rice (Yoshimura & Zaima, 2020), tomatoes (Bednarz, Roloff, & Niehaus, 2019), grape (Berisha et al., 2014) and strawberries (Wang, Yang, Chaurand, & Raghavan, 2021). Applications for animal-based food include raw chicken-meat (E. Marzec, Wojtysiak, Połtowicz, Nowak, & Pedrys, 2016), pork chops (Enomoto, Furukawa, Takeda, Hatta, & Zaima, 2020) and fish (Goto-Inoue, Sato, Morisasa, Igarashi, & Mori, 2019). MS imaging for food analysis has been reviewed with a focus on application examples (Yoshimura et al., 2020) and technical details (Handberg, Chingin, Wang, Dai, & Chen, 2015).

Only very limited data is available for mass spectrometry imaging of processed food. Maslov et al. recently investigated the peptide distribution in dry-cured ham muscle (Rešetar Maslov, Svirkova, Allmaier, Marchetti-Deschamann, & Kraljević Pavelić, 2019). Almost all previously published studies focus on endogenous constituents in food. However, also minor substances such as toxic reaction products, residues, contaminants or food additives are of interest in the context of food safety and authenticity. In the EU, most processed food is subject to regulations regarding maximum levels of residues, contaminants and food additives. In some cases, the spatial distribution is also regulated, e.g. the food additive natamycin. Such components have not been previously analyzed using MS imaging and are described in this work for the first time.

In this study, we want to show the versatility of high-resolution MALDI MS imaging for visualizing the distribution of constituents, ingredients, contaminants and additives. This includes food of both plant and animal origin. We optimized the MS imaging workflow to cater to the specific properties and challenges of the individual samples. Especially for imaging experiments of processed food that consist of multiple ingredients with varying physical and chemical properties, sample preparation procedures differ from established protocols.

We show the distribution of constituents of hardy kiwi, different carrot species (food plants) and German veal sausage (processed meat-based food). We also developed an MS imaging protocol for traditional German gingerbread as an example for highly processed food of plant origin to reveal the spatial distribution of the food contaminant acrylamide. As an example for food additives, we investigated the distribution of the preservative natamycin in cheese, which diffuses from the surface into the cheese. In this case, we have developed a novel analysis approach to investigate the penetration from the surface into the cheese in more detail.

## 2. Methods

### 2.1 Samples

Samples of hardy kiwi (*Actinidia arguta*) as well as orange and purple carrots (*Daucus carota* ssp. *sativus* var. *atrorubens* Alef.) were purchased at a local supermarket.

German veal sausage (“Weißwurst”) was provided by the Max Rubner Institute (Kulmbach, Germany). Samples of Gouda cheese with varying natamycin concentrations (see Table S1 in Supplementary material), the natamycin standard (2.5 % aqueous suspension, Sigma Aldrich, Dreieich, Germany) and acrylamide-contaminated gingerbread (3,200 µg/kg determined by LC-(ESI)-MS/MS) were provided by the Bavarian Health and Food Safety Authority (LGL, Germany).

### 2.2 Sectioning

Frozen samples were sectioned with a cryostat (Leica CM3050, Wetzlar, Germany) at −15 °C - −25 °C object temperature. Section thickness for the food samples were as follows: Kiwi 30 µm, orange carrot 100 µm, purple carrot 50 µm, German veal sausage 16 µm and Gouda cheese varied between 14 µm and 16 µm. Sections were thaw-mounted on adhesion object slides (SuperFrost Plus™, Thermo Scientific™) and stored at −80 °C (gouda cheese at −20 °C) until analysis. Gingerbread-sections of 2 mm thickness were cut off the frozen gingerbread using an electric micro saw and stored at −80 °C until analysis.

### 2.3 MALDI MS imaging

30 mg 2,5-Dihydroxybenzoic acid (DHB, Sigma Aldrich, Dreieich, Germany) were dissolved in 1 ml acetone/water (1:1, Carl Roth, Karlsruhe, Germany) and 0.1 % trifluoroacetic acid (Sigma Aldrich, Dreieich, Germany). 170-200 µL DHB matrix solution was applied to the sections using a pneumatic sprayer system with a nitrogen pressure of 0.7-0.75 bar, a flow rate of 10-15 µL/min and a distance of 10 cm between nozzle and target. Detailed information per sample is given in Table S2 (supplementary material). The section and matrix quality was checked before and after matrix application with a digital microscope (Keyence VHX-5000, Osaka, Japan).

Mass spectrometric data were acquired with an AP-SMALDI10 ion source (TransMIT GmbH, Giessen, Germany) coupled to a Q-Exactive-HF Orbitrap mass spectrometer (Thermo Scientific, Bremen, Germany). The MALDI source is equipped with a 60 Hz Nitrogen laser (λ = 337 nm) and one scan event consisted of 30 laser pulses. The laser was focused down to a spot size of 10 µm and the energy on the target varied between 0.5 µJ and 2.0 µJ. For the gingerbread samples, analyses were performed with 12.5 µJ laser energy and 60 laser shots per pixel.

All imaging measurements were performed in positive ion mode with a mass resolution of 240,000 at *m/z* 200 (120,000 at *m/z* 200 for German veal sausage sample). Matrix clusters were used as lock masses for the appropriate mass range to gain a mass accuracy typically better than 1.5 ppm (Treu & Römpp, 2021). The *m/z* ranges varied for each application and are given in Table S3 in the supplementary material as well as raster and pixel size for the corresponding imaging experiments.

On Gouda sections MS/MS experiments were also performed. The details are provided in the supplementary material Figure S7 and Figure S8.

### 2.4 Data analysis

MS data were analyzed with the QualBrowser of the Thermo Xcalibur 4.0 software. Tentative compound identification was based on accurate mass unless stated otherwise. MS imaging data were converted to the open file format imzML (Schramm et al., 2012) using ‘jimzML Converter’ (Version 2.0.4) (Alan M. Race, Styles, & Bunch, 2012) and ‘imzML Validator’(A. M. Race & Römpp, 2018).

MSiReader 1.0 (Bokhart, Nazari, Garrard, & Muddiman, 2018) and Mirion (Paschke et al., 2013) (version 3.2.64.12) were used to generate MS images with a selected *m*/ window of +/− 2.5 ppm. Preprocessing steps (such as normalization) are indicated in the corresponding figure caption. Detailed penetration analysis of natamycin was performed using our newly developed semiautomatic penetration tool based on SpectralAnalysis (A. M. Race, Palmer, Dexter, Steven, Styles, & Bunch, 2016) and written in MATLAB (R2016b), which consists of two parts: i) Lipid based edge detection to determine the interface ii) calculation of penetration plots. More details are given in the result section (Figure 5) and in the supplementary material (Figure S10).

**Figure 5:**
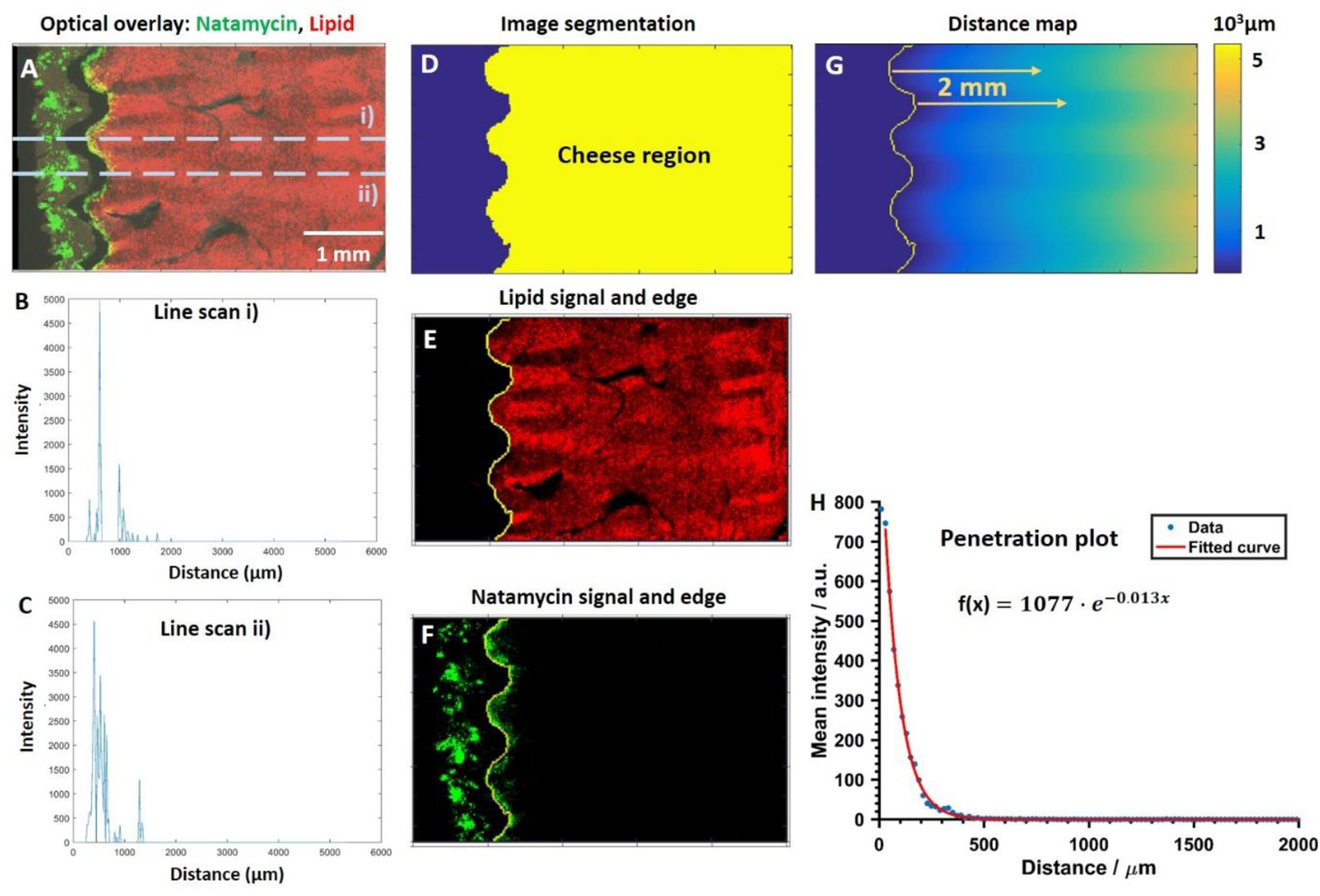
Description of developed method for determining the cheese edge and then calculating a ‘penetration plot’ for natamycin into the cheese. A RG-overlay composite – R lipid SM(d34:1) ([M+Na]^+^, m/z 725.55680), G natamycin ([M+Na]^+^, m/z 688.29396), optical image. B+C line scans, showing the intensity of natamycin against the lateral distance in µm. D: image segmentation E: lipid signal and calculated edge from D. F: natamycin signal displayed with edge from D. G: distance map from edge of cheese surface. H: penetration plot of natamycin into cheese including exponential fit. The mean intensity of the natamycin signal is given for a certain distance from the surface of the cheese against the distance in x-direction.

## 3. Results and Discussion

In the following, we demonstrate the power of applying mass spectrometry imaging to food science applications. The examples range from ingredients, contaminants and additives in samples as diverse as plant and animal-based, fresh and processed food. Motivation and specific experimental protocols for each application are discussed in the respective section.

### 3.1 Constituents in fresh plant-based food

#### 3.1.1. Hardy kiwi

The hardy kiwi (*Actinidia arguta*) is an essentially hairless grape-sized fruit with edible peel and an intense sweet and sour taste reminding of tropical fruit (Hallett & Sutherland, 2005) (Fisk, McDaniel, Strik, & Zhao, 2006). Apart from its nutritional value, it contains several health-promoting bioactive constituents, such as vitamin C, polyunsaturated fatty acids and polyphenols (Kim, Beppu, & Kataoka, 2009; Nishiyama, Yamashita, Yamanaka, Shimohashi, Fukuda, & Oota, 2004; Park et al., 2011). One half of a 30 µm thin cross section of a hardy kiwi was analyzed in positive ion mode with a step size of 45 µm and a mass range of *m/z* 250-1000 (Fig 1A). The mass spectral signals shown in Fig 1B were acquired from a single pixel. The MS images in Figure 1C show the distributions of the lipid glyceryl trilinolenate (TG(54:9), [M+K]^+^, *m/z* 911.65255, red) dihexoses ([M+K]^+^, *m/z* 381.07937, green) and anthocyanin quercetin ([M+H]^+^, *m/z* 303.04993, blue). The combination of these single ion images (RGB MS image) and the optical image (Figure 1D) shows that the structure of the sample was retained throughout sample preparation and data acquisition. The distribution of the imaged constituents can be directly linked to structures in the hardy kiwi. These compounds – and all other compounds reported in this study - were identified by accurate mass, i.e. mass accuracy was better than 1.5 ppm. This high specificity of the mass spectral data is particularly important in mass spectrometry imaging measurements as the complexity of the (food) sample cannot be reduced by chromatographic separation (Römpp et al., 2013). The [M+K]^+^-adduct of the lipid TG(54:9) (*m/z* 911.65255), for example, was detected in the single MS spectrum in Figure 1B with a mass deviation of 0.26 ppm. The mass accuracy for the whole measurement can be determined as the root mean square error (RMSE) of the mass deviation for each individual spectrum containing the targeted ion. The calculated RMSE for all pixels in this RGB MS image were 1.14 ppm (3427 spectra), 0.80 ppm (32705 spectra) and 0.55 ppm (127821 spectra) for the signals of glyceryl trilinolenate (*m/z* 911.65255, red), quercetin (*m/z* 303.04993, blue) and dihexose (*m/z* 381.07937, green), respectively.

This allows for reliable compound identification and an image generation with a bin width of +/−2.5 ppm. This provides specific information on the distribution of analytes and significantly reduces the risk of interference by neighboring peaks. As expected, dihexoses (Figure 1D, green) were found with high intensities in the pericarp of the hardy kiwi. Glyceryl trilinoleate (Figure 1D, red) is the most abundant triglyceride in kiwi seed oil (Piombo et al., 2006), which supports our assignment. Due to the different cutting planes of the seeds, the relative abundance of the lipid varies between the two seeds in the measured section. Kiwi fruit contain an average of 2660 mg/100 g total phenolic content and the peel in particular is rich in polyphenols ((Baranowska-Wójcik & Szwajgier, 2019; Kim et al., 2009). This is consistent with the high intensities of quercetin (Figure 1C, blue), found here in the peel. Similar distributions of additional anthocyanins such as cyanidin and pelargodinin are shown in Figure S1 (supplementary material). The example of the hardy kiwi shows that MS imaging provides the combination of specific molecular information with spatial information, which can be used to link the distributions of components to specific functions or metabolic processes. This information can be obtained for a wide range of compound classes with different physicochemical properties as shown in the following examples.

#### 3.1.2. Carrots

Carrots are popular root vegetables and valued for their high content of beta-carotene (approx. 7.6 mg/100 g) (Souci, Fachmann, & Kraut, 2008). A carrot has a very solid consistency and is hard to section in a frozen state. Therefore, a 100 µm thick cross section of an orange-colored carrot was analyzed in positive ion mode with a step size of 50 µm and a mass range of *m/z* 100-1500. Using MALDI MS imaging it was possible to detect beta-carotene ([M]^+^, *m/z* 536.43820). The MS-image is shown with the corresponding optical images of the carrot in Figure 2A. The single pixel mass spectrum of beta-carotene is shown next to the MS image in Figure 2A and a mass deviation of 0.52 ppm was determined for the shown signal. The RMSE for the whole measurement is 1.29 ppm (35981 spectra). Carotenes are highly nonpolar and therefore difficult to detect in MALDI experiments due to their low ionization efficiency. However, the beta-carotene distribution is visibly comparable to the orange-colored regions in the optical image and thus confirms our results. Apart from the well-known orange-colored variety, differently colored carrots are also cultivated, such as yellow or purple varieties. Purple carrots (*Daucus carota* ssp. *sativus* var. *atrorubens* Alef.) have an attractive purple-colored outer core and cortex due to high concentrations of anthocyanins. We could detect these anthocyanins in an MS imaging measurement and reconstruct the purple ring in the cross section as an MS image of cyanidin ([M]^+^, *m/z* 287.05557, RMSE: 1.02 ppm (2100 spectra)). The MS image is shown with the corresponding optical image in Figure 2B.

**Figure 2:**
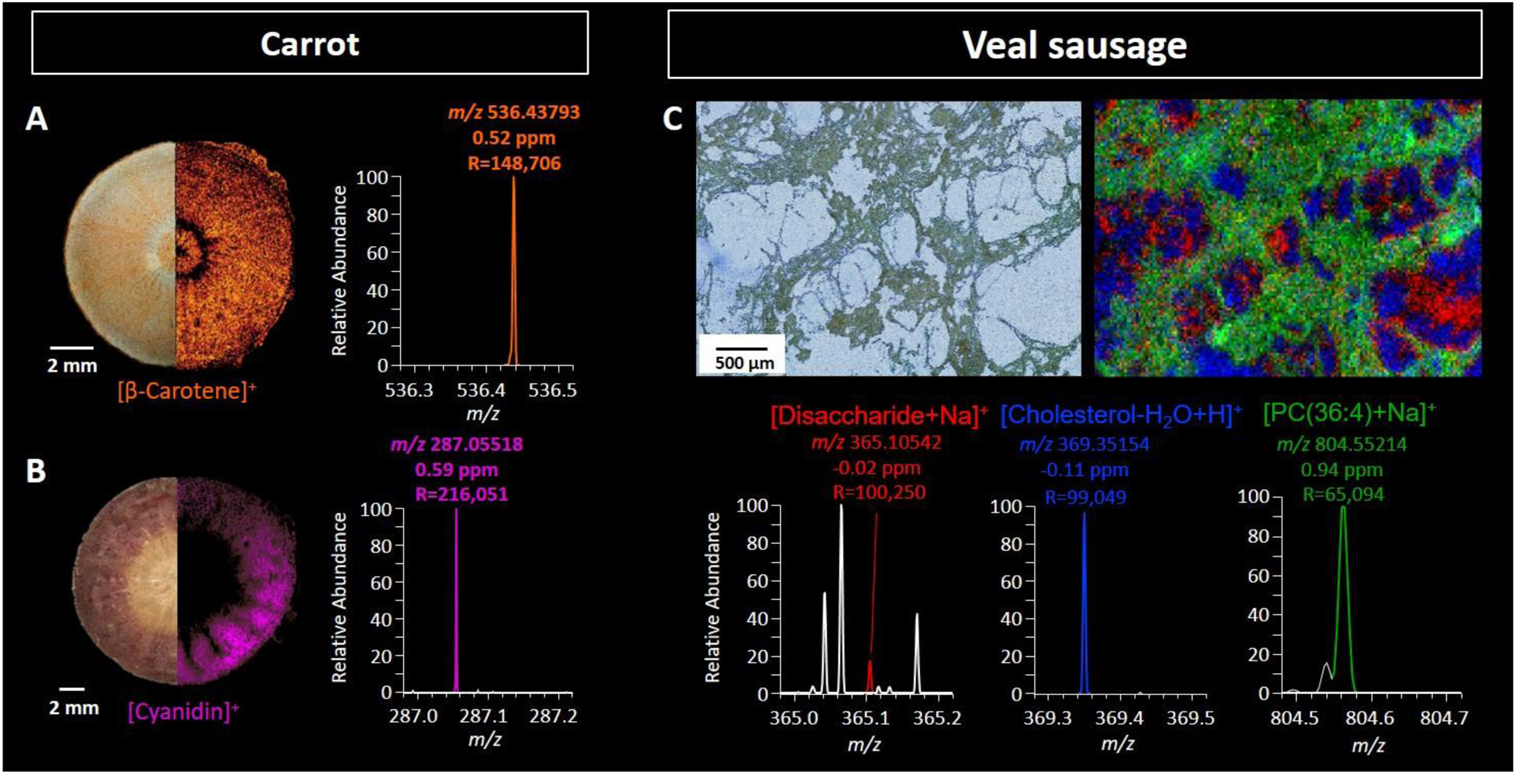
MS imaging of constituents in carrot species and veal sausage: A: orange carrot: Juxtaposition of optical image and single ion image of beta-Carotene ([M]^+^, m/z 536.43765, orange), pixel size 50 µm. B) Purple carrot: Juxtaposition of optical image and single ion image of Cyanidin ([M]^+^, m/z 287.05501, purple), pixel size 80 µm. C: Optical image and RGB MS image of three constituents of veal sausage: Disaccharide ([M+Na]^+^, m/z 365.10544, red), PC(36:4) ([M+Na]^+^, m/z 804.55138, green) and Cholesterol ([M-H2O+H]^+^, m/z 369.35158, blue), pixel size 20 µm.

A previous study used Raman mapping to gain insight into compartment-specific differences in carotenogenesis in orange- and purple-colored carrots (Baranska, Baranski, Schulz, & Nothnagel, 2006). They were able to differentiate three different carotenoids based on Raman spectroscopy. However, their assessment of anthocyanins (as a compound class) was based only on visual color perception (Baranska et al., 2006). In contrast, MS imaging can provide information not only on the spatial distribution, but also on specific molecule identities for a wide range of compound classes.

### 3.2 Ingredients in animal-based processed food

Using MS imaging, it is not only possible to investigate constituents in fresh, but also in processed food. Homogenizing a mixture of ingredients is a common method in food processing, resulting in dramatically altered physicochemical properties that have to be taken into account for MS imaging analysis – especially for sample preparation. German veal sausage is produced by mincing trimmed rosé veal with a low content of fat and tendons, high-fat pork and bacon with the optional addition of herbs and spices such as parsley, onion, pepper and mace (Lebensmittelbuch-Kommission, 2019). Consequently, regions of low fat-content can be found in direct proximity to high-fat regions and shreds of connective tissue. These high-fat regions influenced the sample preparation process. Sectioning was performed at −25 °C and matrix application was optimized towards a ‘wetter’ spray (higher flow rate; see Table S2 in the supplementary material). Using this adapted MS imaging workflow, the low-fat and high-fat regions can be clearly distinguished by visualizing water-soluble and fat-soluble constituents as shown in Figure 2C. In this example, the *m/z* of water-soluble disaccharide ([M+ Na]^+^, *m/z* 365.10544, RMSE: 0.68 ppm (24394 spectra), red) and the fat-soluble cholesterol ([M– H_2_O + H]^+^, *m/z* 369.35158, RMSE: 0.62 ppm (27491 spectra), blue) show nearly complementary distributions. The lipid PC(36:4) ([M+Na]^+^, m/z 804.55138, RMSE: 1.23 ppm (30549 spectra), green), a phosphatidylcholine naturally occurring in biological membranes, covers the entire tissue. The 16 µm thin section in Figure 2C was imaged with a pixel size of 20 µm in the mass range of *m/z* 200-900. Single ion images of all constituents shown in the RGB overlay of Figure 2C are provided in the supplementary material (Figure S2). In addition, ingredients of plant origin could be localized by tracing the *m/z* of chlorophyll-derivatives originating from added herbs, as shown in Figure S3 (supplementary material). This shows that not only optically visible structures can be reproduced by MS imaging also in processed food, but also invisible tissue regions can be distinguished by tracing suitable marker compounds.

**Figure 3:**
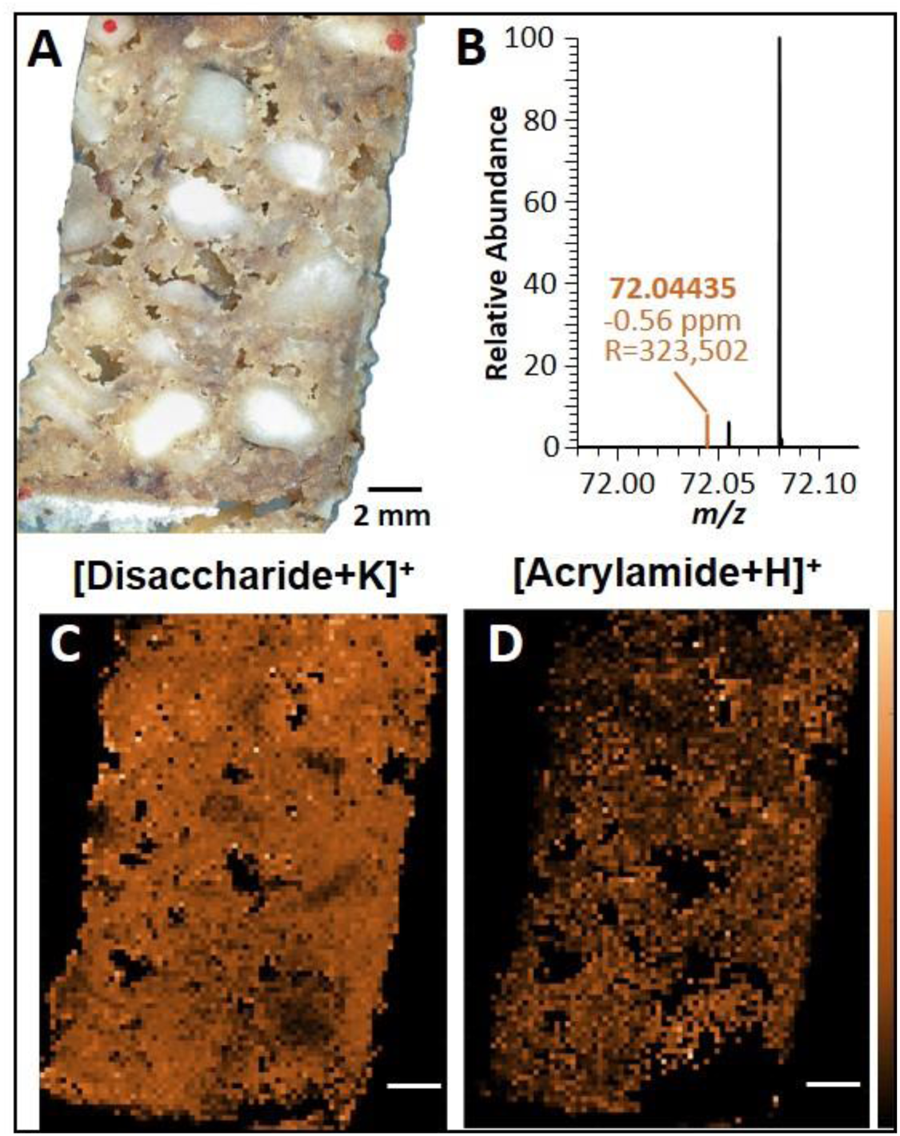
MS imaging of German gingerbread. A: optical image, the red dots represent reference points for the MALDI laser B: Single pixel mass spectrum of acrylamide C: MS image of Disaccharide [M+K]^+^, m/z 381.07937, mass range m/z 300-900, pixel size 200 µm D: MS image of process contaminant acrylamide [M+H]^+^, m/z 72.04439, mass range m/z 50-150, pixel size 200 µm. MS images are TIC-normalized.

### 3.3. Contaminants in plant-based processed food

In addition to constituents shown in the previous examples, the distributions of minor components, such as contaminants, are of interest in food analysis. Food contaminants are legally defined as substances that are unintentionally added to food during the production chain, which includes primary production, preparation and packaging. A prominent example is the carcinogen acrylamide, which is formed from the naturally occurring constituents asparagine and sugars, when prepared at low moisture and temperatures higher than 120 °C (Stadler et al., 2002). The occurrence of acrylamide in food is continuously under discussion, especially since the new Regulation (EU) 2017/2158, establishing mitigation measures and benchmark levels, entered into force^1^. Figure 3 presents results of a newly developed MS imaging workflow for the investigation of the acrylamide distribution in traditional German gingerbread. Due to its very dry and hard consistency, the standard cryosectioning procedure was not applicable for the gingerbread samples. Instead, we developed a sectioning protocol using an electric micro saw to obtain sections of approx. 2000 µm thickness.

The solvent system was adapted to a higher proportion of acetone (see Table S2, supplementary material) to gain a better matrix crystallization on the gingerbread sections. The thickness and texture of the sections led to an uneven/rough surface which can influence the ionization efficiency during the MALDI process while rastering across the sample with the laser. We addressed this problem by using reference points (red marker pen) on different spots across the sample to choose the appropriate/average distance between sample and laser optics. Two alternating scan intervals of *m/z* 50-150 and *m/z* 300-900, respectively, were set to cover a large *m/z* range with high mass resolution and optimized ion intensities for all analytes of interest.

In Figure 3A, a gingerbread section is shown which was prepared and measured with the described optimized MS imaging workflow at 200 µm pixel size. Our results show nearly ubiquitous presence of disaccharides throughout the sample surface, demonstrated in Figure 3C by the potassium adduct ([M+K]^+^, *m/z* 381.07937, RMSE: 0.44 ppm (3995 spectra)), indicating the success of the established workflow. Furthermore, sufficient intensity for the spatially resolved detection of acrylamide ([M+H]^+^, *m/z* 72.04439, RMSE: 1.05 ppm (2335 spectra) Figure 3B) was achieved. The contaminant is distributed throughout all gingerbread ingredients, i.e. it could be detected within the dough as well as the wafer and nuts (Figure 3D). The successful elucidation of the acrylamide distribution in German gingerbread is the first MS imaging study of a contaminant in processed food and also demonstrates the analytical capability of MS imaging for the detection of low-abundant food components.

### 3.4 Additives in animal-based processed food

In contrast to contaminants, food additives are intentionally added during the production process to obtain certain properties such as color or flavor of the processed food. In some cases, not only a concentration limit is defined for food additives, but the location of a regulated component in the sample is also specified. A prominent example is the antifungal drug natamycin, which is approved as a preservative in the European Union (E235). According to Regulation (EC) No 1333/2008, natamycin may be added to meat products as well as hard and semi-hard cheeses^2^. Gouda is a prominent example for a semi-hard cheese with dry-matter between 49 % and 57 % and a minimal maturation of 5 weeks^3^. Apart from the maximum level of 1 mg/dm^2^ on the cheese surface, the regulation states a penetration limit of 5 mm for natamycin^4^. Official food analyses in Germany are listed in the Official Collection of Methods according to § 64 of the German Foodstuffs and Feed Code (LFGB)^5^. According to the official method, natamycin is quantified by HPLC-DAD after methanolic extraction from a cheese slice close to the surface and from the bulk (see supplementary material Figure S4).

**Figure 4:**
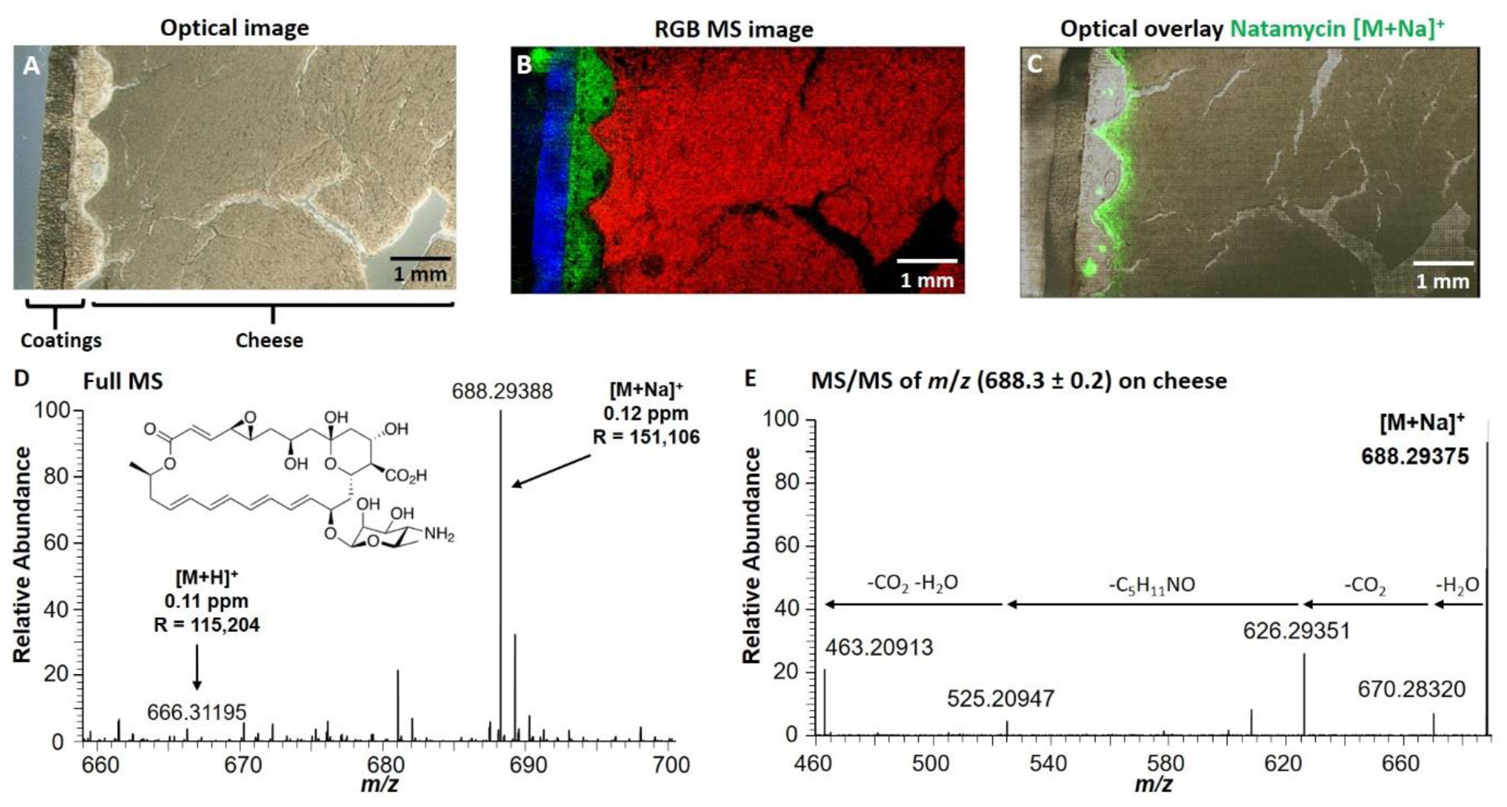
MS imaging of Gouda cheese A: Optical image of a Gouda cheese section. B: RGB MS image of characteristic signals for the coatings m/z 257.22641 (blue) and m/z 320.243103 (green) and the lipid SM(d34:1) ([M+Na]+, m/z 725.55680, red). MS/MS spectra of this lipid can be found in Figure S7, supplementary material. MS imaging measurement was acquired with a pixel size of 20 µm C: Overlay of natamycin signal ([M+Na]+, green) and optical image. 4D: Single MS spectrum acquired from a natamycin hotspot on the Gouda cheese, [M+H]+ at m/z 666.31202 and [M+Na]+ at m/z 688.29396 are highlighted with their corresponding mass deviation in ppm and mass resolution R (FWHM). E: Mean MS/MS spectrum of [M+Na]+ (isolation window: m/z 688.3 ± 0.2) at NCE = 25.

This method allows quantification, but does not provide spatial information. In cooperation with the *Bavarian health and food authority* (LGL), we have therefore developed a MALDI MS imaging workflow to investigate the natamycin penetration into cheese sections. A schematic description of the workflow can be found in Figure S5 (supplementary material). Special care must be taken while sectioning the cheese samples to avoid cross-contamination of natamycin from the surface towards the cheese bulk. The sectioning process of cheese in general is very challenging, due to its high lipid content and varying textures between different cheeses. Furthermore, the presence of holes in certain cheese varieties can easily cause cracks during sectioning. The high lipid content would normally lead to lower cutting temperatures, but the texture of certain Goudas was too brittle if the temperatures were too low. This caused problems while sectioning and made thaw-mounting on glass slides impossible. Therefore, the cutting temperature needed to be optimized separately for every cheese sample. Analogous to our previous study on drug compound imaging in mouse model tissue, the glass slides were warmed from behind by finger contact in the area of the section in order to ensure proper sample mounting (Treu, Kokesch-Himmelreich, Walter, Holscher, & Römpp, 2020). In the optical image of a Gouda section (16 µm thickness) in Figure 4A, two layers of coating and the cheese region can be seen. The MS imaging experiment shown in Figure 3B was conducted with a pixel size of 20 µm in the mass range of *m/z* 200-800. We were able to detect characteristic compounds for the three different sample regions, e.g. two coating compounds in blue and green, and one lipid in the cheese bulk in red (see Figure S6 for single ion images). The lipid could be identified as SM(d34:1) ([M+Na]^+^ *m/z* 725.55680, RMSE 0.58 ppm (55198 spectra)), see also supplementary material, Figure S7. The RGB MS image matches the optical image very well, which shows that the imaging workflow was successfully developed and retains the structure of the sample.

**Figure 6:**
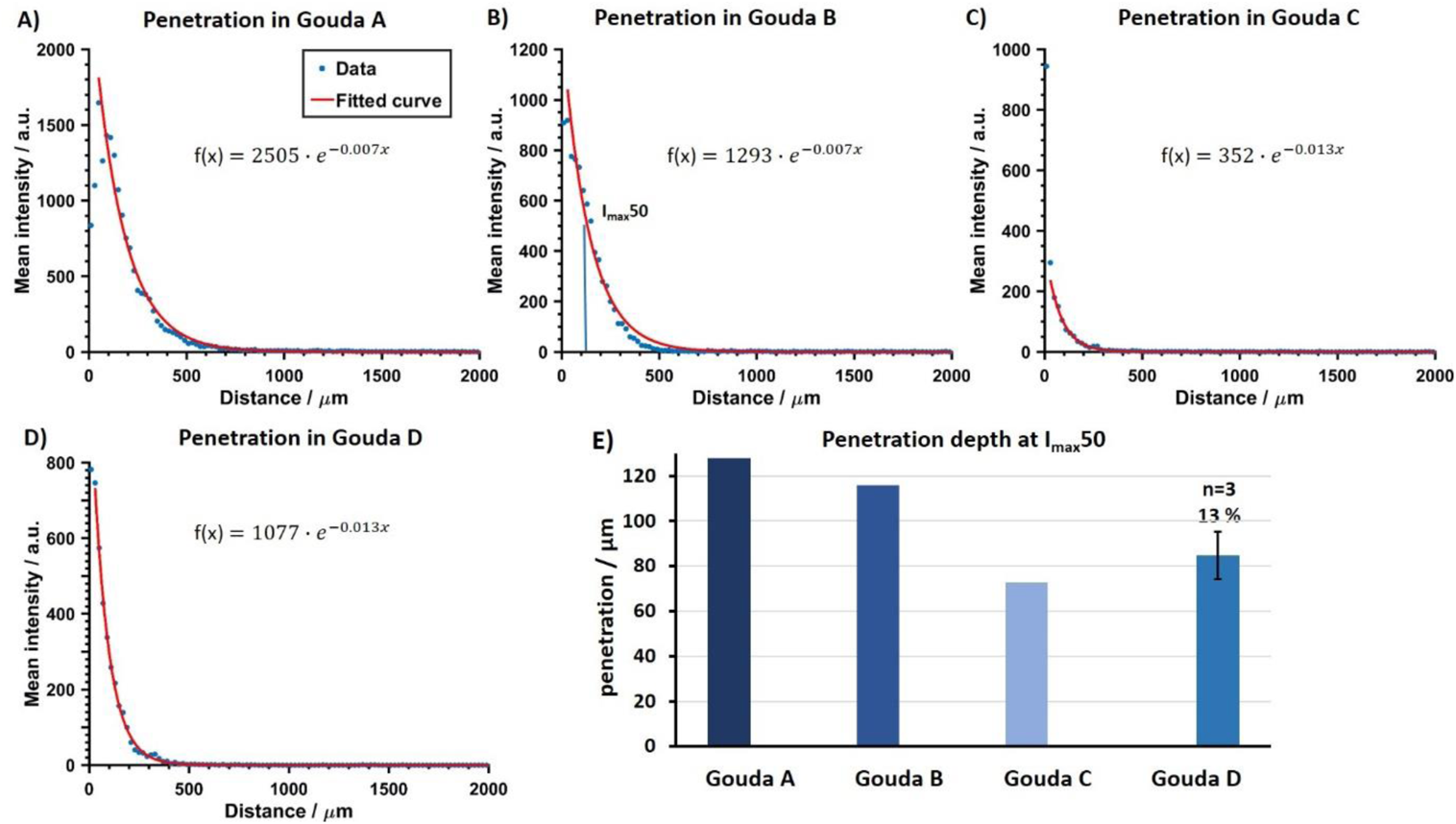
Comparison of natamycin penetration in Gouda A, B, C, D. A-D: Penetration plots for different Gouda samples. Mean intensity against distance from the surface in µm. E: Comparison of penetration depth at 50% of maximum intensity (I_max50_). For Gouda D the mean of three neighboring sections with standard deviation is shown, relative error 13%.

Furthermore, using MS imaging it is possible to visualize the distribution of natamycin, which is not visible in the optical image. The distribution of the sodiated natamycin adduct ([M+Na]^+^, *m/z* 688.29396) is shown in Figure 4C as an overlay with the optical image. It can be observed that natamycin occurs in this sample mainly in the rind of the cheese. The sodium adduct ([M+Na]^+^) showed the highest intensity in comparison to the protonated ion ([M+H]^+^) and the potassium adduct ([M+K]^+^) in all of our measurements. This can be explained by the industrial application of natamycin on cheese as an aqueous sodium solution and the natural occurrence of sodium in cheese.

In Figure 4 D, a single pixel mass spectrum from the measurement in Figure 4C is shown in the mass range of *m/z* 650 – 700. The [M+H]^+^ signal (*m/z* 666.31202) and the [M+Na]^+^ adduct (*m/z* 688.29396) was detected with a mass deviation of 0.11 ppm and 0.12 ppm, respectively. Over the whole measurement, a mass accuracy ≤ 1 ppm could be obtained (RMSE: 0.94 ppm in 1190 spectra and 0.63 ppm in 5199 spectra, respectively). For additional confirmation, MS/MS spectra of the [M+Na]^+^ adduct in cheese were acquired (Figure 4E). Four characteristic product ions were detected, which match the MS/MS spectra of the pure natamycin standard (Figure S8, supplementary material) and thus confirm the identity of natamycin in the Gouda sample.

In addition, a Gouda sample with no declared natamycin – confirmed by the HPLC-DAD reference method - was imaged. In the corresponding MS image of this sample no natamycin signal was present (Figure S9, supplementary material). This combination of accurate mass detection, on-sample MS/MS fragmentation and verification with a blank sample as shown here by the example of natamycin provides for an additional level of certainty for the identification of food components. Using the MS imaging approach, we could confirm the presence of natamycin in multiple cheese samples. Different spatial distributions of natamycin were observed in each sample. One example is shown in Figure 5A, where the sodiated natamycin signal ([M+Na]^+^ *m/z* 688.29396) is shown in green and the lipid SM(d34:1) (corresponding to the lipid in Figure. 4B) is shown in red in an optical overlay. Here, natamycin was detected not only in the rind, but also in the coating of the cheese. Application methods for natamycin onto the cheese surface vary between producers; it can be applied onto the cheese surface before the coating or premixed with the coating, explaining the observed differences.

The determination of the penetration of an analyte in the MS image is not straightforward. A simple approach to determine the penetration of natamycin into the cheese would be the use of line scans. Two examples are shown in Figure 5B+C; the intensity of natamycin is plotted against the x-direction of the image (given in µm). Comparing the two line scans it becomes clear that a single line scan is not representative for the whole cheese section. Due to the rough surface of the cheese, and the noisy line scans, it is difficult to determine the edge of the cheese bulk and the exact depth of penetration, which strongly influences data interpretation.

Similar problems have been described in the literature in case of a drug penetration studies in human skin (Bonnel et al., 2018) and 3D cell cultures (Machalkova et al., 2019). Bonnel *et al* used a semi-automated workflow to correlate the drug concentration to a certain depth in the skin based on a user-defined region of interest. Machalkova *et al*. evaluated penetration depths of the investigated drug in the 3D cell culture with the MALDI MS imaging supported by complementary information from laser scanning confocal microscopy. In contrast, we developed a data analysis approach that solely relies on MS data and includes an automated identification of the region of interest (interface between cheese and coating). This workflow is briefly explained in the following and more details can be found in the supplementary material, Figure S10.

As a first step, we use the lipid signal, shown in Figure 5A, which occurred in all our investigated cheese samples to generate a mask for the cheese (Figure 5D). For generating this mask, image-processing techniques were used to remove noise and fill in gaps to be able to determine a smooth cheese edge (Figure 5E). Therefore, it is possible to differentiate between natamycin pixels on cheese and off cheese (Figure 5F). Natamycin signals in the coating (‘off cheese’) are not considered for the following analysis since they are not in the cheese.

After edge detection, the lateral distance to the edge is calculated for every pixel in the cheese region and can be plotted as a ‘distance map’ (Figure 5G). The mean intensity of the natamycin signal is then calculated for all pixels with the same lateral distance from the edge. The mean intensity values were plotted against the distance in *µm* to generate a penetration plot, shown in Figure 5H. The zero value on the x-axis represents the cheese surface (the edge in the distance map). Data points close to the edge show the highest natamycin intensity, with the intensity decreasing with higher distances from the edge. This trend can be expressed in an exponential function and the fit equation describes the diffusion of natamycin into the cheese bulk. Every step in the data analysis workflow and the corresponding figures and graphs are automatically generated in one run by the developed data analysis tool.

Additional Gouda samples with different natamycin concentrations from different producers (Table S1, supplementary material) were analyzed in the same way. The penetration plots of four samples are depicted in Figure 6A-D. Sample Gouda D corresponds to the penetration plot shown in Figure 5H. The penetration plots show that natamycin diffused into the cheese in all cases, however, different penetration behaviors could be observed for the four examples. To compare the four cheese samples, penetration depths at 50% of the maximum intensity value (I_max50_) were calculated using the fit equations. The comparison of these I_max50_ values is shown in Figure 6E.

In order to assess the reproducibility of our approach, two additional (serial) sections from the same cheese were measured and analyzed in case of Gouda D. All three penetration plots of the neighbouring sections show the same trend and the penetration depths are comparable (Figure S11, supplementary material). In Figure 6E, the mean value of (85 ± 11 µm) is shown for these three replicates of Gouda D. The relative standard deviation of 13% indicates that our approach can be applied with reasonable reproducibility to determine the penetration depths. The I_max50_ values can be used to compare the diffusion behaviour of natamycin in different samples. Gouda A shows the highest value, and Gouda C the lowest. The penetration analysis shows that none of the investigated cheese samples exceeded the penetration limit of 5 mm of the applicable EU regulation. Apart from the penetration analysis, our approach allows comparing mean intensities in a defined area of multiple sections and samples. A comparison to the HPLC-DAD results from the first 2 mm of the cheese, which were measured by LGL, is given in Figure S12 (supplementary material). The MS imaging results follow the same trend as the homogenate analysis from the HPLC-DAD reference method, while providing additional spatial information as discussed above.

With our MS imaging workflow, we were able to determine the spatial distribution of natamycin in cheese samples. This constitutes the first spatially resolved detection of a food additive in food. The advantage is that we can investigate the penetration of natamycin in much more detail compared to the routine LC analysis (which has a “spatial resolution” of 5 mm). With our advanced data analysis approach, it was possible to compare the natamcyin penetration between different samples. This penetration is most likely dependant on the application method for natamycin, which varies between brushing, dipping, spraying and addition to the brine (Kammerlehner, 2015). Therefore, our approach could contribute towards less exposure to natamycin for the customers, which would be in line with Article 11 of Regulation (EC) No 1333/2008 stating that for food additives “the level of use shall be set at the lowest level necessary to achieve the desired effect”^6^.

## 4. Conclusions

MS imaging can be used for a wide range of food-related applications. Our results show that the unique combination of molecular and spatial information can provide the distribution of different compound classes in various food sample matrices. We were able to detect highly nonpolar compounds such as triglycerides Tin the hardy kiwi and beta-carotene in the orange carrot. We could also visualize the distribution of very polar compounds such as disaccharides and anthocyanins in hardy kiwi and purple carrot. Our data include not only fresh plant-based food, but also processed food comprising meat, dairy and bakery products. The specific properties of each sample required the development of a dedicated workflow. In the case of processed food, the workflow is more challenging due to the fragile and heterogeneous texture of the samples as they consist of multiple ingredients with varying physical and chemical properties. The properties of the processed food samples in our study ranged from lipid rich (veal sausage/cheese) to very dry (gingerbread). We successfully visualized different distributions of food constituents (disaccharides, lipids), even in homogenized samples such as the German veal sausage.

In addition to constituents, we also investigated minor components of processed food, i.e. a food additive and a contaminant. Our investigations of an acrylamide in gingerbread and natamycin in cheese are the first MS imaging analyses of a contaminant and a food additive distribution in processed food. Natamycin was detected in varying distributions between the investigated cheese samples. With this example, we also showed that MS imaging goes beyond mere visualization of compounds, and can provide additional information on the penetration behavior of food additives. The developed data analysis approach can be used as a general tool to investigate diffusion processes by MS imaging in a wide range of other applications. In conclusion, our results show that MS imaging has great potential to complement established methods in food analysis by providing deeper information about spatial distributions of food components, in particular those underlying official regulations.

## Acknowledgements

We would like to thank Holger Zorn (Justus Liebig University Giessen) for initial discussions on the topic of natamycin MS imaging and Jan Lauter for first experiments on sample preparation. We would also like to thank Holger Knapp and Jennifer Mels from the *Bavarian Health and Food Safety Authority* (LGL, Erlangen, Germany) for providing the gingerbread samples and the Max Rubner-Institute (MRI, Kulmbach, Germany) for providing the German veal sausage.

## Founding sources

This work was supported by the Deutsche Forschungsgemeinschaft (DFG) INST 91/373-1-FUGG and the TechnologieAllianzOberfranken (TAO).

## Appendix A. Supplementary data

The Supplementary Material provides further details on samples, sample preparation and MALDI imaging experiment settings. Additional MS images for the hardy kiwi and veal sausage are shown. For the cheese analysis, detailed information on the LC-experiment workflow, the MALDI imaging workflow and the data analysis workflow are presented. More details on MS/MS data, blank cheese data and reproducibility are provided.

## Supplementary material

### 1. Material and Methods

**Table S1:** Gouda samples provided by the LGL (Bavarian health and food safety authority). Shown natamycin concentrations were measured by the LGL using the HPLC-DAD reference method. All listed samples were also investigated in this study by MS imaging.

| Sample | Natamycin declaration | mg/dm<sup>2</sup> Surface | mg/dm<sup>2</sup> Bulk |
| --- | --- | --- | --- |
| Gouda Blank | no | < 0.01 | < 0.01 |
| Gouda A | yes | 0.57 | < 0.01 |
| Gouda B | yes | 0.42 | < 0.01 |
| Gouda C | yes | 0.11 | < 0.01 |
| Gouda D | yes | 0.38 | < 0.01 |

**Table S2:** Sample preparation: Detailed information of sectioning and spraying parameters for all samples in this study. *The ginger bread sample was stored at −80 °C before sectioning, but it was not sectioned in a cryotome, please see the “method section” in the main paper.

| Sectioning |  |  | Spray Parameters |  |  |  |
| --- | --- | --- | --- | --- | --- | --- |
| Sample | Temperature/°C | thickness/μm | DHB Solution | Flow rate/μl/min | gas pressure/bar | Volume/mL |
| Hardy Kiwi | -24 | 30 | Aceton/H<sub>2</sub>O (50:50) + 0.1% TFA | 10 | 0.7 | 175 |
| Orange carrot | -15 | 100 | Aceton/H<sub>2</sub>O (50:50) + 0.1% TFA | 10 | 0.7 | 175 |
| Purple carrot | -24 | 50 | Aceton/H<sub>2</sub>O (50:50) + 0.1% TFA | 10 | 0.7 | 175 |
| Veal sausage | -25 | 16 | Aceton/H<sub>2</sub>O (50:50) + 0.1% TFA | 15 | 0.75 | 170 |
| Gouda | -20 | 14/16 | Aceton/H<sub>2</sub>O (50:50) + 0.1% TFA | 10 | 0.7 | 170 |
| Ginger bread | -80* | 2000 | Aceton/H<sub>2</sub>O (70:30) + 0.1% TFA | 10 | 0.7 | 200 |

**Table S3:** Detailed data acquisition parameters for MALDI MS imaging. The ginger bread section was measured in an alternating scanning mode (Treu, Kokesch-Himmelreich, Walter, Holscher, & Römpp, 2020) with a step size of 100 µm x 200 µm (XxY). Every second scan event was measured with one of the given mass ranges (m/z 50-150 or m/z 300-900). This leads to a pixel size of 200 µm in the MS images.

| Sample | Raster size | Step size | mass range |
| --- | --- | --- | --- |
| Hardy Kiwi | 320x540 | 45 | 250-1000 |
| Orange carrot | 237x237 | 50 | 100-1500 |
| Purple carrot | 244x216 | 80 | 200-800 |
| Veal sausage | 240x170 | 20 | 200-900 |
| Gouda A | 340x185 | 20 | 200-800 |
| Gouda B | 310x145 | 20 | 200-800 |
| Gouda C | 330x175 | 20 | 200-800 |
| Gouda D | 315x150 | 20 | 200-800 |
| Gouda blank | 285x235 | 20 | 200-800 |
| Ginger bread | 76x97 | 100x200 | 50-150 / 300-900 |

### 2. Results and discussion

#### Constituents in fresh plant-based food

**Figure S1:**
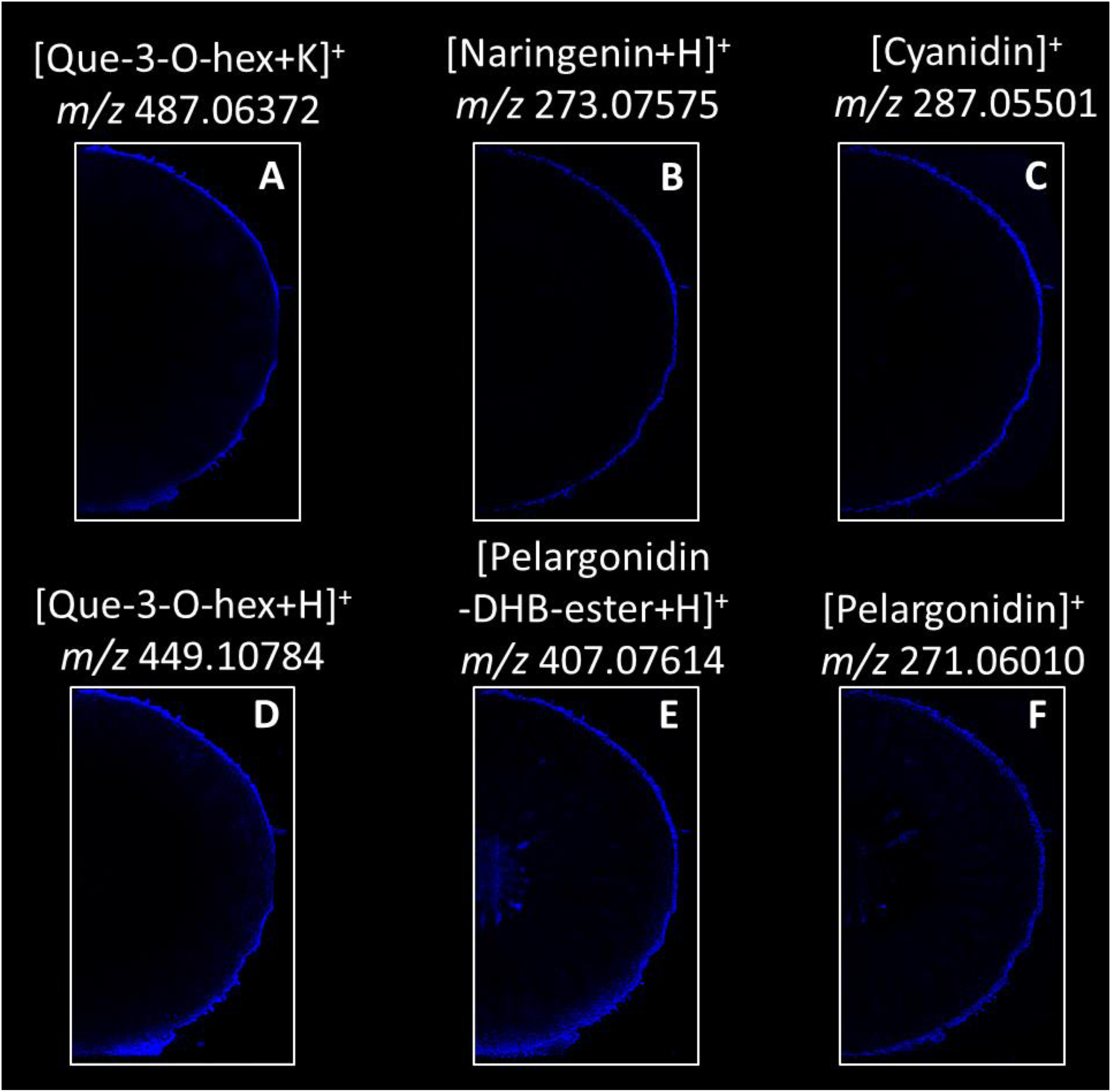
Distributions of additional anthocyanin signals in hardy kiwi (Figure 1). All depicted analytes show high intensities in the peel, pelargonidin is also detectable in the stem (E,F).

#### Ingredients in animal-based processed food

**Figure S2:**
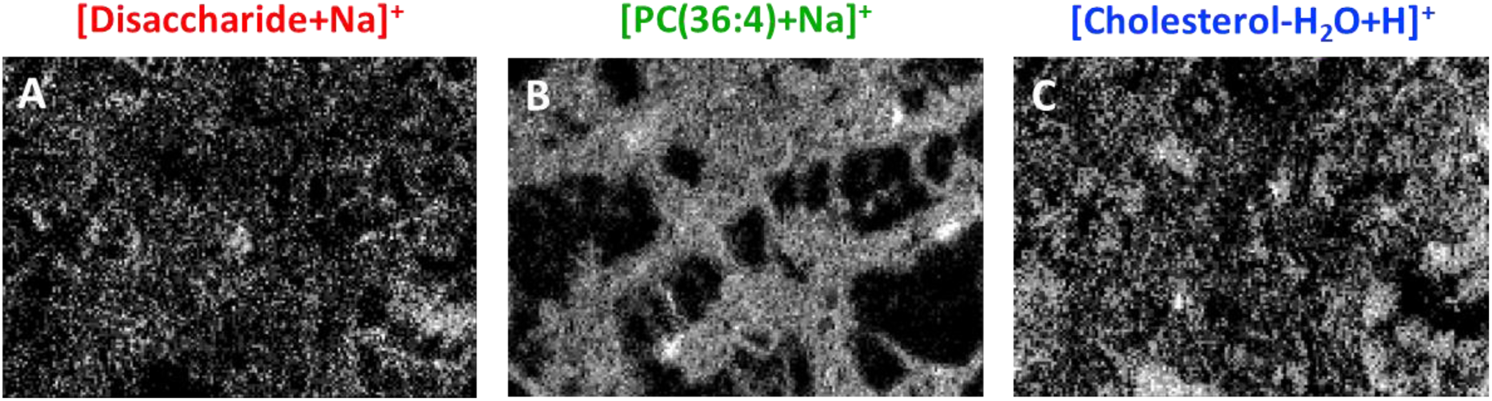
Single ion images for the RGB MS image of three constituents of German veal sausage in Figure 2C. A: Disaccharide ([M+Na]^+^, m/z 365.10544), B:PC(36:4) ([M+Na]^+^, m/z 804.55138), and C: Cholesterol ([M-H2O+H]^+^, m/z 369.35158).

**Figure S3:**
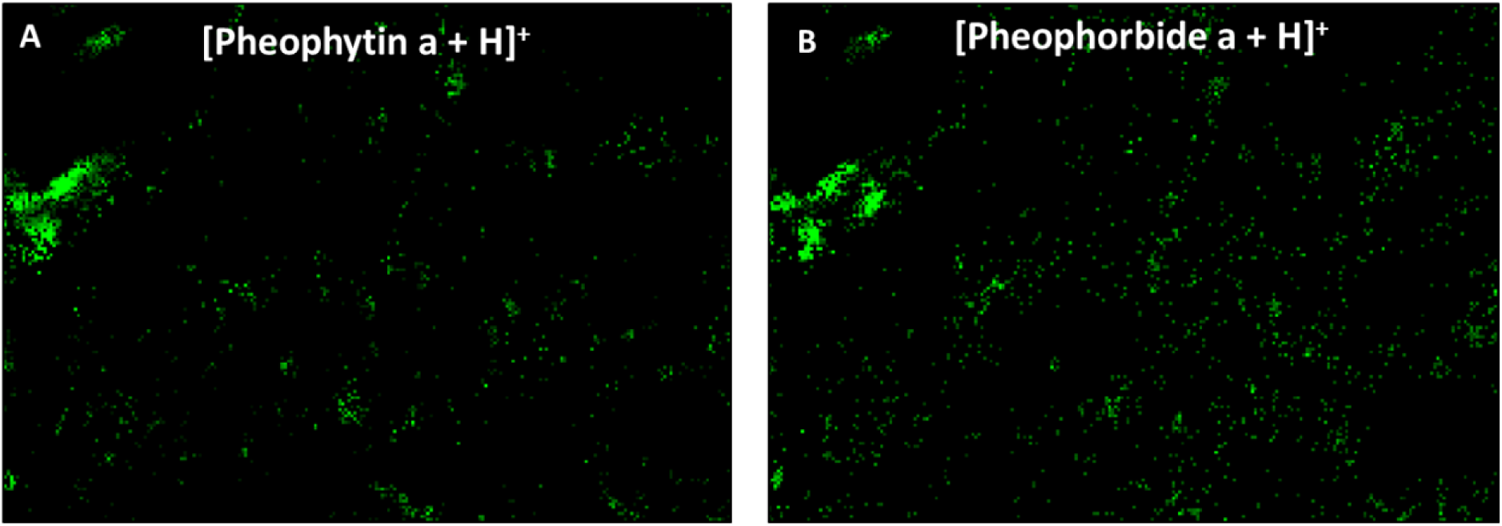
Distribution of chlorophyll-derivatives originating from added herbs in German veal sausage in Figure 2C. A: Pheophytin ([M+H]^+^, m/z 871.57319) and B: Pheophorbide ([M+H]^+^, m/z 593.27584).

#### Additives in animal-based processed food

**Figure S4:**
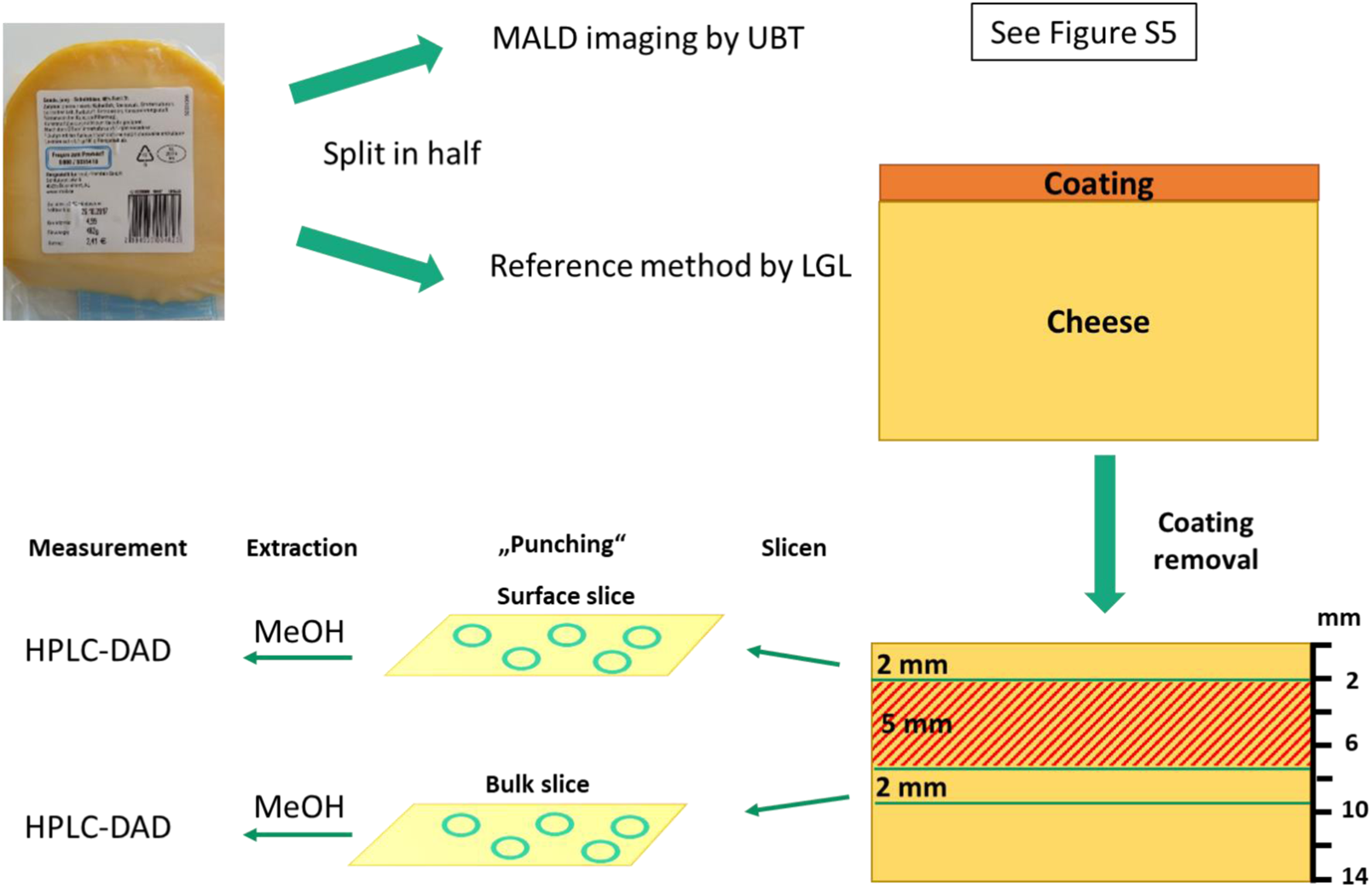
LGL-DAD reference method for quantification of natamycin: Adapted from § 64 LFGB - German Food and Feed Code.

The cheese samples were split in half. One half was shipped on ice to the University of Bayreuth for MALDI imaging analysis. The other half was investigated using the LC-DAD reference method by the LGL. After removing the coating of the cheese, a slice of 2 mm was set aside for further analysis, another 5 mm were discarded and another slice of 2 mm was used also for further analysis. The first slice was collected to determine the surface concentration and the second as reference for the bulk concentration. From both slices 5 tablets were punched out and subjected to methanol extraction followed by HPLC-DAD analysis.

**Figure S5:**
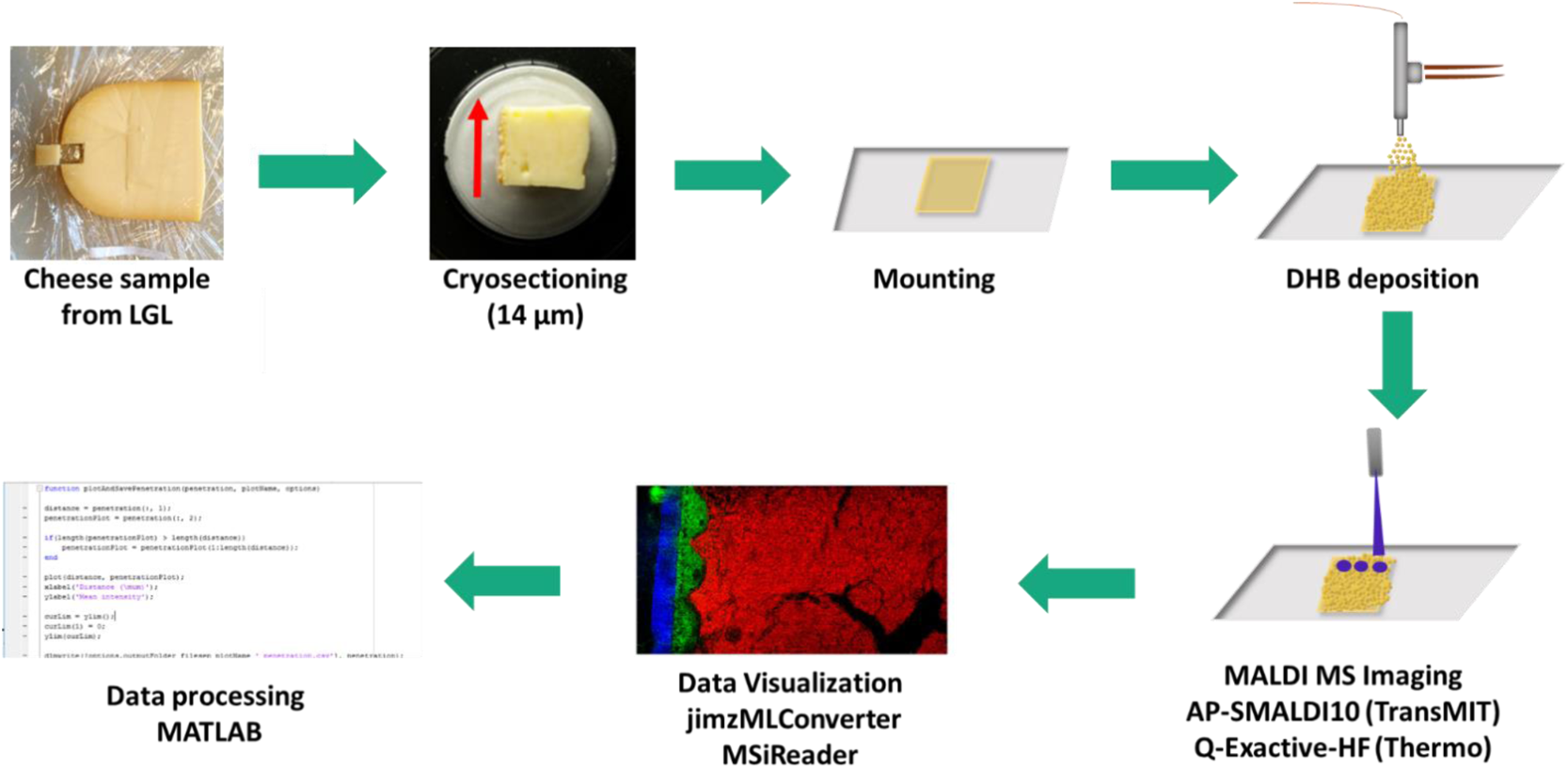
M*A*LDI *MS imaging workflow for all cheese samples.* Small pieces were cut out of cooled Gouda samples and were stored at −20°C until use. For cryo-sectioning, the Gouda pieces were attached to a sample holder of a cryostat (CM3050 S, Leica, Germany) with distilled water only. In accordance to the maturity level and the fat content, the Gouda pieces were sectioned at different chamber (CT) and object (OT) temperatures (CT in the range of −19°C to −21°C and the OT in the range of −19°C and −23°C). The sections (14 and 16 µm thickness) were thaw-mounted on adhesion object slides (SuperFrost Plus™, Thermo Scientific™) by warming the glass slides from behind by finger contact in the area of the section in order to ensure proper sample mounting (Treu, Kokesch-Himmelreich, Walter, Holscher, & Römpp, 2020). The sections were stored at −20°C until analysis. Prior to matrix application the sections were dehydrated in a vacuum desiccator. DHB was applied using a pneumatic sprayer. All data were acquired using the AP-SMALDI10 high-resolution MALDI imaging ion source (TransMIT GmbH) which was coupled to a Q-Exactive HF Orbitrap mass spectrometer (Thermo Fisher). MS imaging data was converted to the open file format imzML (Schramm, Hester, Klinkert, Both, Heeren, Brunelle, et al., 2012) using the ‘jimzML converter’ (Version 2.0.4) (Alan M. Race, Styles, & Bunch, 2012) and the ‘imzML validator’ (A. M. Race & Römpp, 2018). MSiReader 1.0 (Bokhart, Nazari, Garrard, & Muddiman, 2018) was used to generate MS images and m/z windows of 2.5 ppm were chosen to generate all shown MS images. Detailed penetration analysis of natamycin was performed using our newly developed semiautomatic penetration tool based on SpectralAnalysis (A. M. Race, Palmer, Dexter, Steven, Styles, & Bunch, 2016) and MATLAB (Version 3.2.64.12) which consists of two parts: i) Lipid based edge detection to determine the interface ii) calculation of penetration plots.

**Figure S6:**
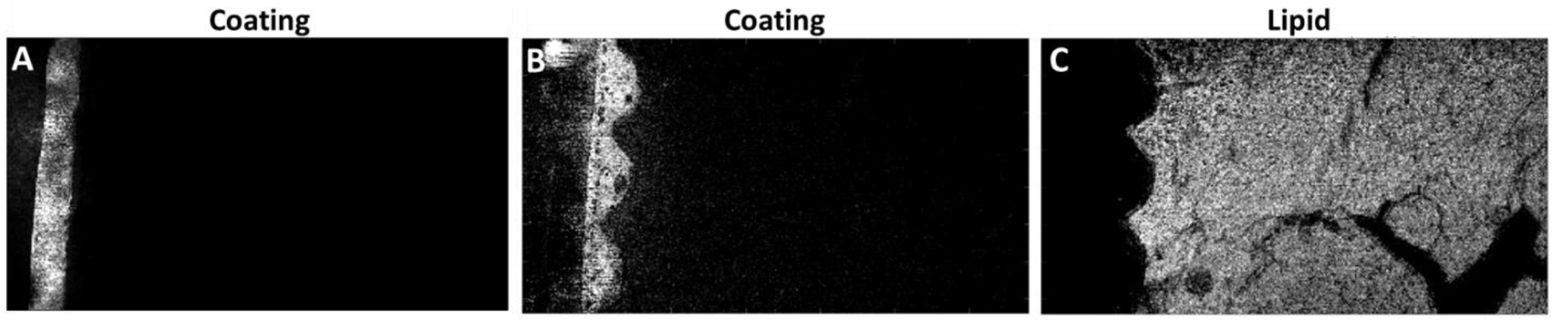
Single ion images of the RGB MS image in Figure 4B: Characteristic compounds for the coatings m/z 257.22641 in A, m/z 320.24290 in B and the lipid SM(d34:1) ([M+Na]^+^, m/z 725.55680) in C.

**Figure S7:**
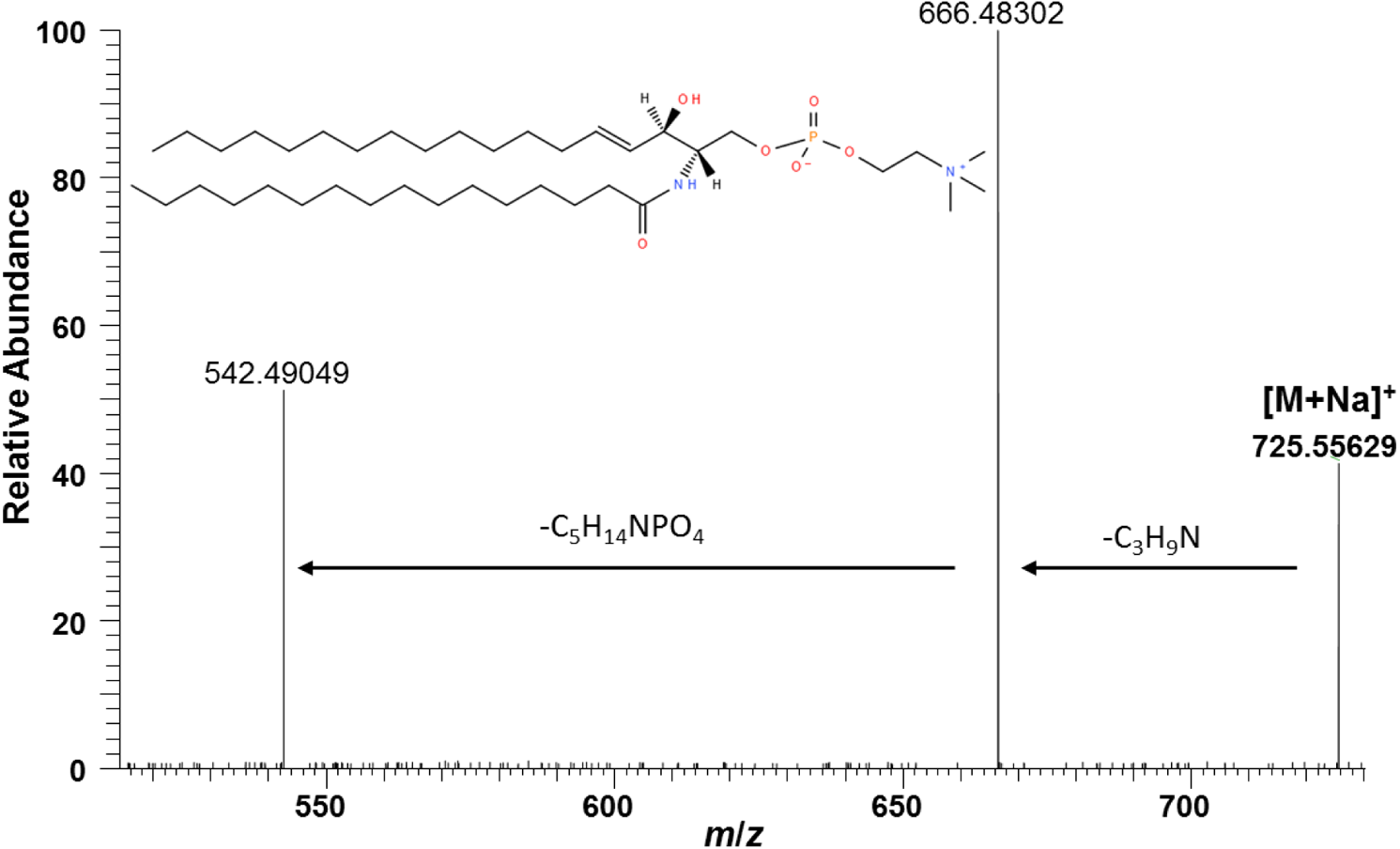
Average MS/MS spectrum of lipid signal m/z 725.55680 in Figure 4B (HCE = 25, isolation window = m/z ± 0.2). The two characteristic neutral losses of C_3_H_9_N and C_5_H_14_NPO_4_ confirm the lipid species sphingomyelin. Therefore we annotated the signal with the database result SM(d34:1) (lipidmaps.org).

**Figure S8:**
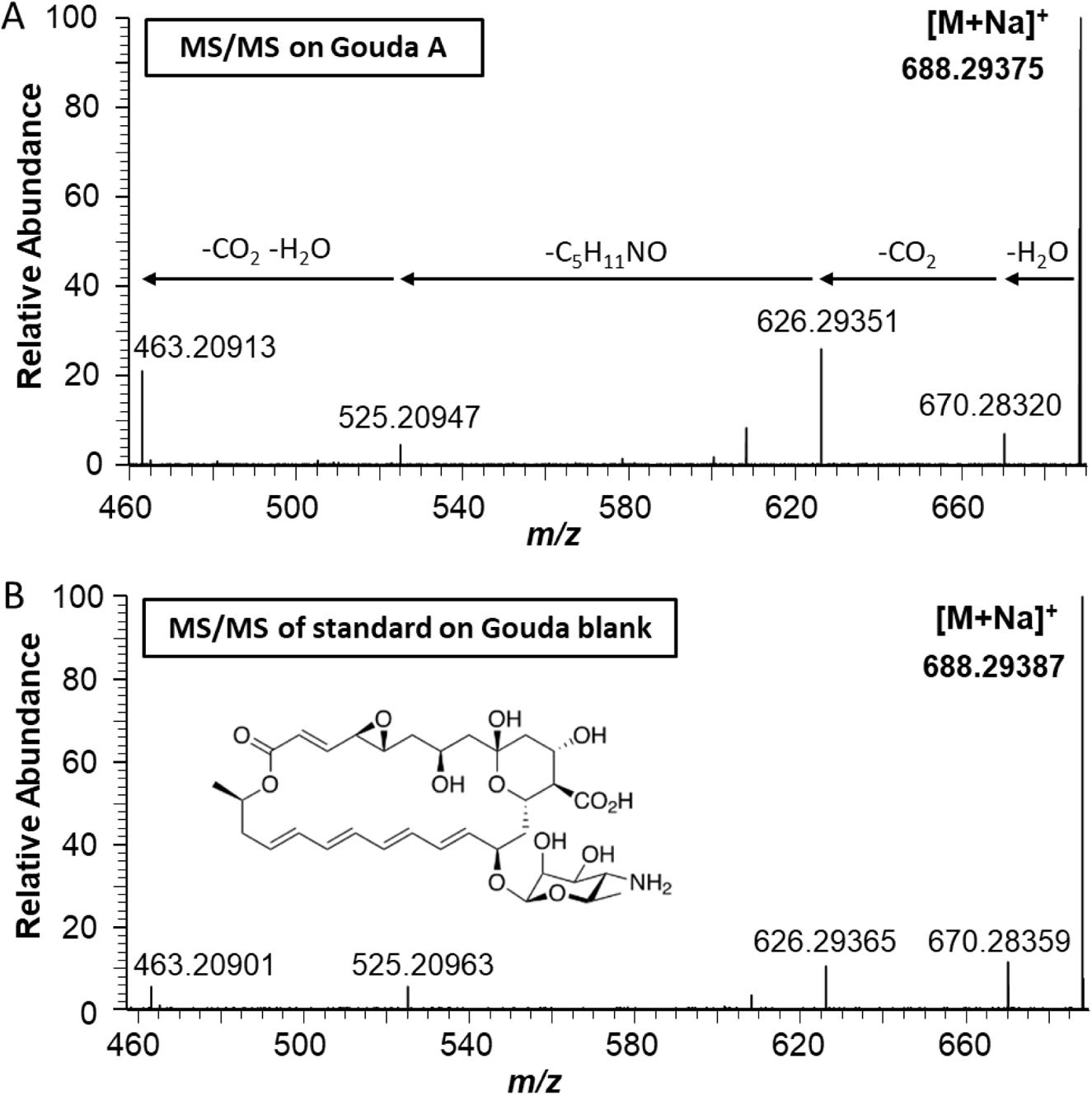
Comparison of MS/MS spectra of natamycin: A: MS/MS of natamycin hot spot on Gouda sample A (HCE = 25, isolation window = m/z ± 0.2). B: MS/MS of natamycin standard. A 2.5 % aqueous natamycin suspension was diluted in methanol/water (50:50, v/v) and pipetted on a Gouda blank section. DHB was applied as for all other cheese samples. For MS/MS experiments an isolation window of m/z ± 0.2 and a collision energy of 20 was used. The same fragment ions could be detected with slightly different intensities, which can be explained by the different collision energies. In combination with the high mass accuracy in the full MS spectrum (see Figure 4) the signal at m/z 668 can be unequivocally identified as the sodiated natamycin.

**Figure S9:**
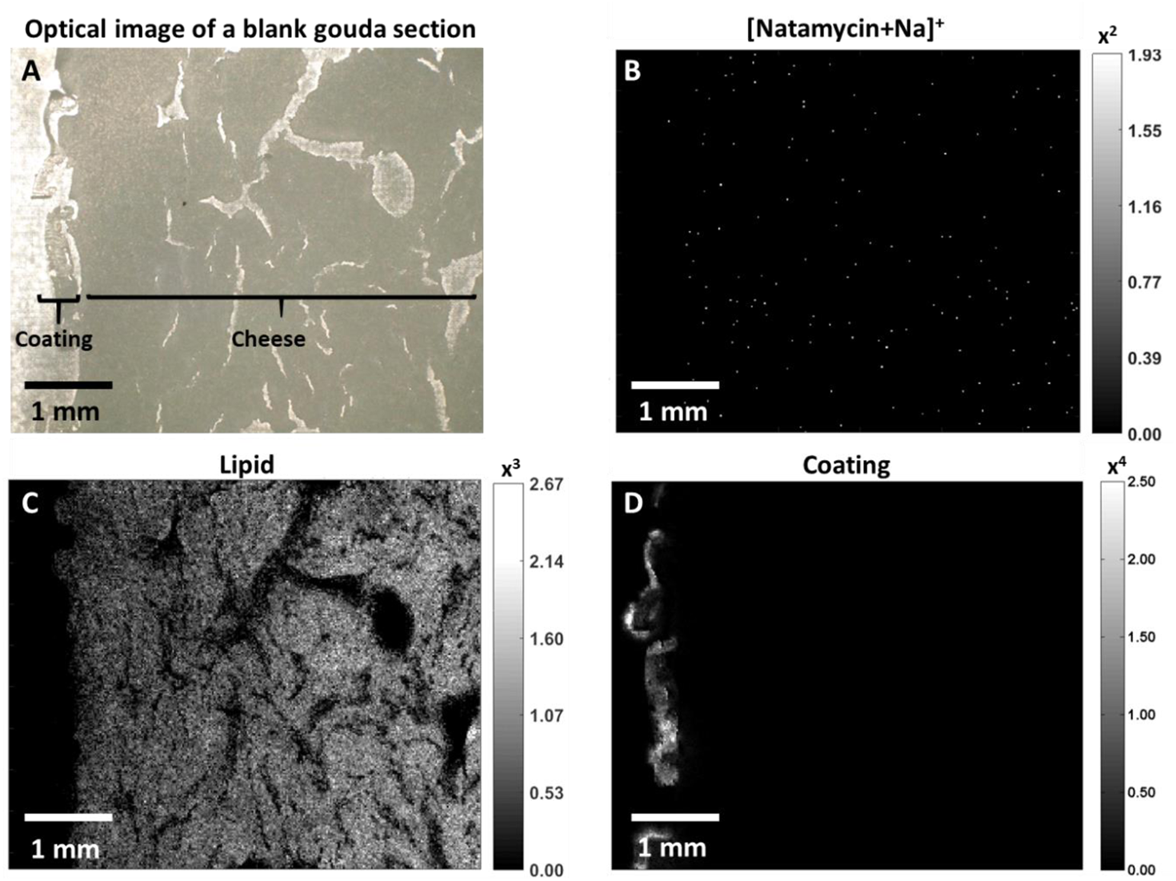
MS imaging of blank Gouda section: A) optical image, B) Natamycin signal ([M+Na]^+^, m/z 688.29396, C) Distribution of lipid SM(d34:1) ([M+Na]^+^, m/z 725.55660), D) coating signal m/z 512.40041. The lipid and coating distribution match the optical image well. No natamycin could be detected in this Gouda section.

#### Penetration analysis

**Figure S10:**
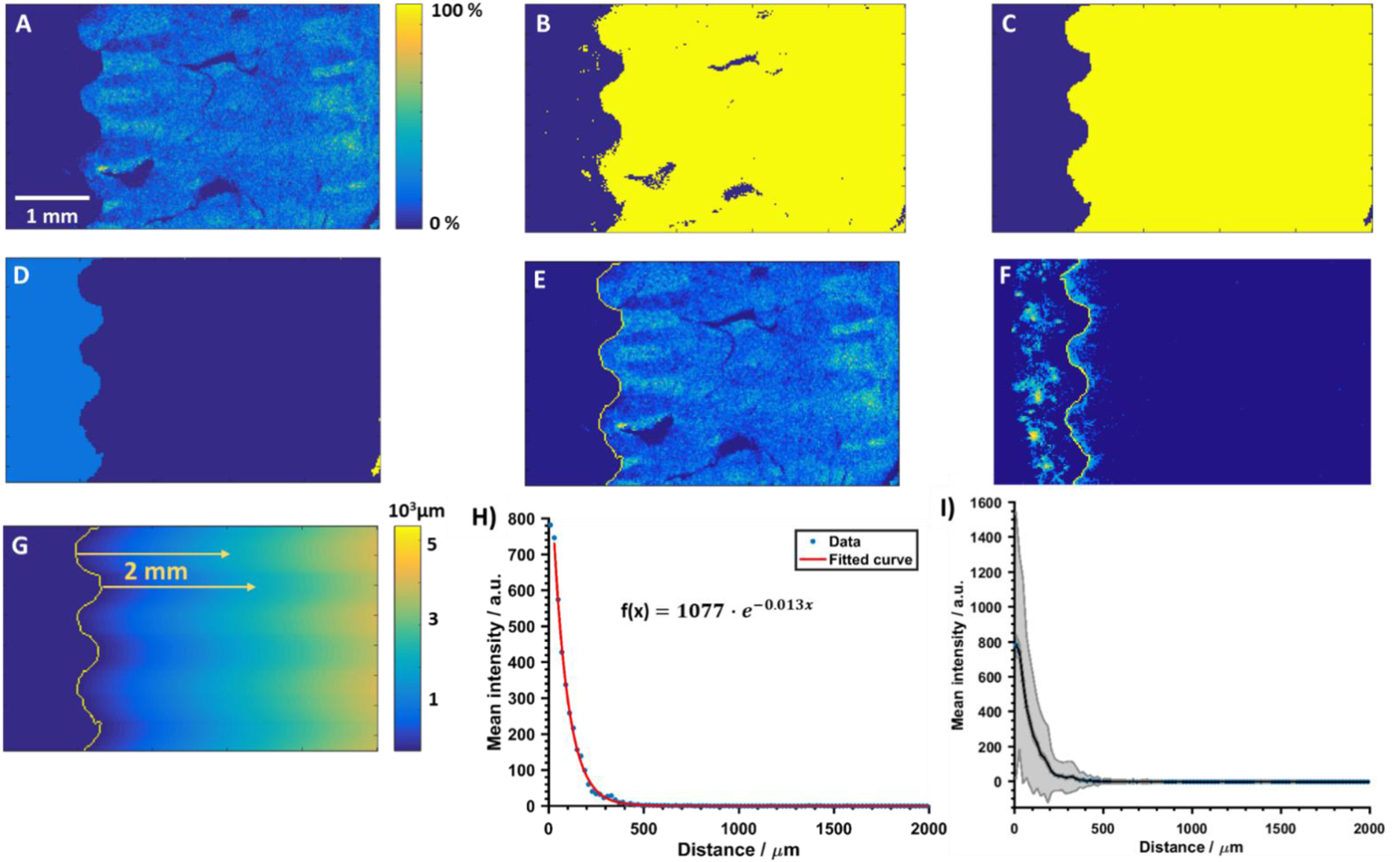
D*e*tailed *developed data analysis approach.* An ion image representing a lipid signal which is only present in the cheese region was generated (Fig S10A). The ion image was then thresholded to form a binary mask (Figure S10B). Small features were removed using the mathematical morphology opening (erosion followed by dilation) with a disk-shaped structuring element (radius = 3) leaving a binary mask of the cheese bulk. Holes in the binary image were then filled, with the largest continuous remaining area forming the cheese mask (Figure 10C). The inverse of this mask formed the ‘off cheese’ mask (Figure S10D). The cheese boundary was defined as the edge between the cheese and off-cheese masks and is shown with the lipid signal in Figure S10E and with the natamycin signal in Figure S10F. Using the cheese boundary, it is possible to calculate the distance (Euclidean distance in the ‘x’ direction) from the cheese edge of every pixel within the cheese region, shown pictorially in the distance map in Figure (S10G). The intensity of the analyte of interest at every pixel within the cheese can be combined with the distance information (binned at the pixel size of 20 um) and was averaged (mean intensity) for a certain distance to form a penetration curve (Figure S10H). Each data point represents the mean intensity of natamcyin for all pixels with this certain distance. An exponential function was then fit to this data and the standard error calculated (shown in shaded area in Figure S10I). Figure S10I shows that the first pixels have the highest error. As the first pixel in the distance plot is at the cheese edge, it is possible that this covers both ‘off’ and ‘on’ cheese regions, and so was omitted from the exponential curve fitting process.

**Figure S11:**
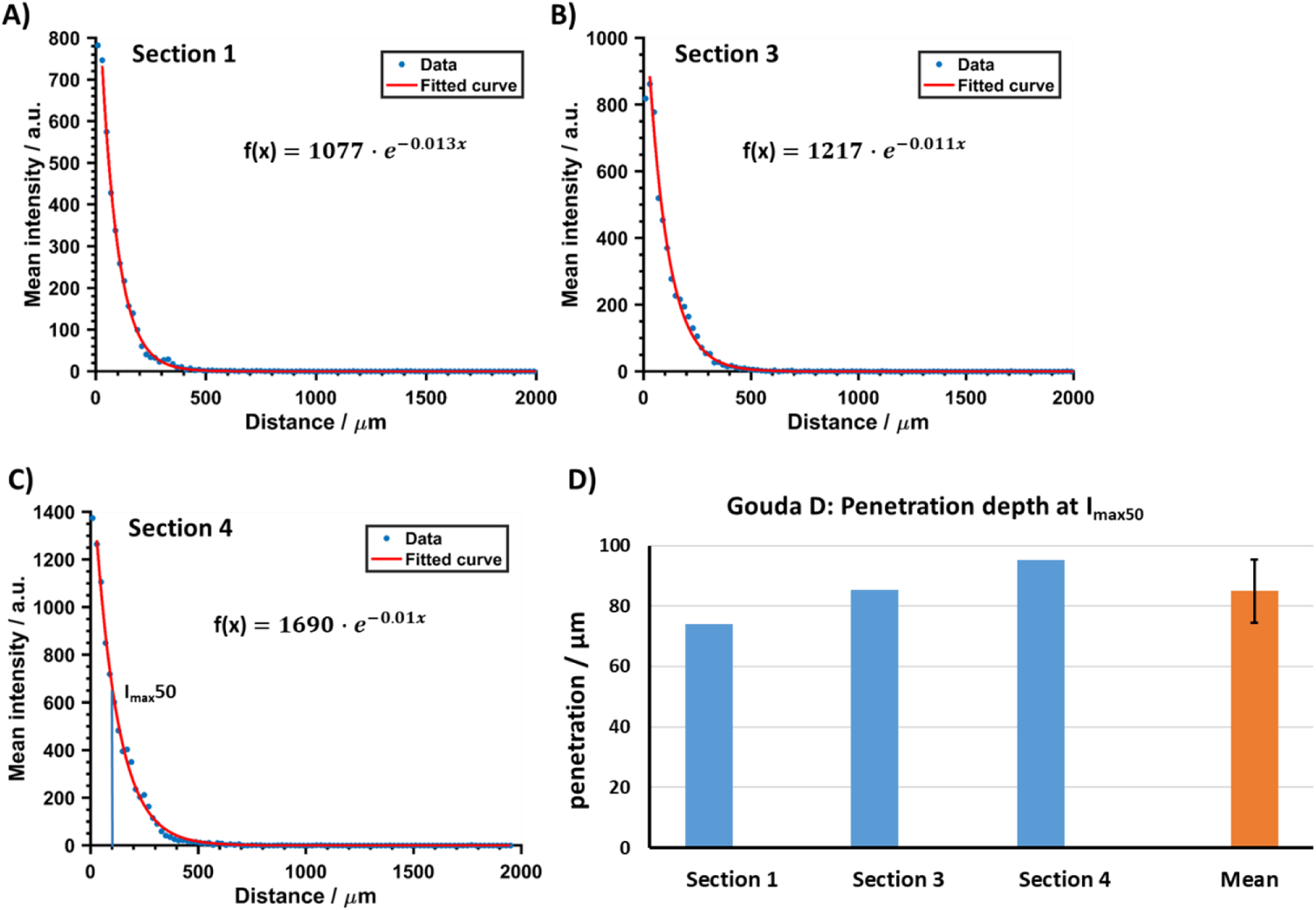
Penetration curves for three neighboring sections of Gouda D. A) Penetration curve including exponential fit and equation of section 1 (same as Figure 5H and 6D). B) Penetration curve including exponential fit and equation of section 3 of the same cheese. C) Penetration curve including exponential fit and equation of section 4 of the same cheese. D: Comparison of calculated I_max_50 values and calculated mean and standard deviation for all three sections. A relative error of 13% was calculated and shows the good reproducibility of the whole workflow.

**Figure S12:**
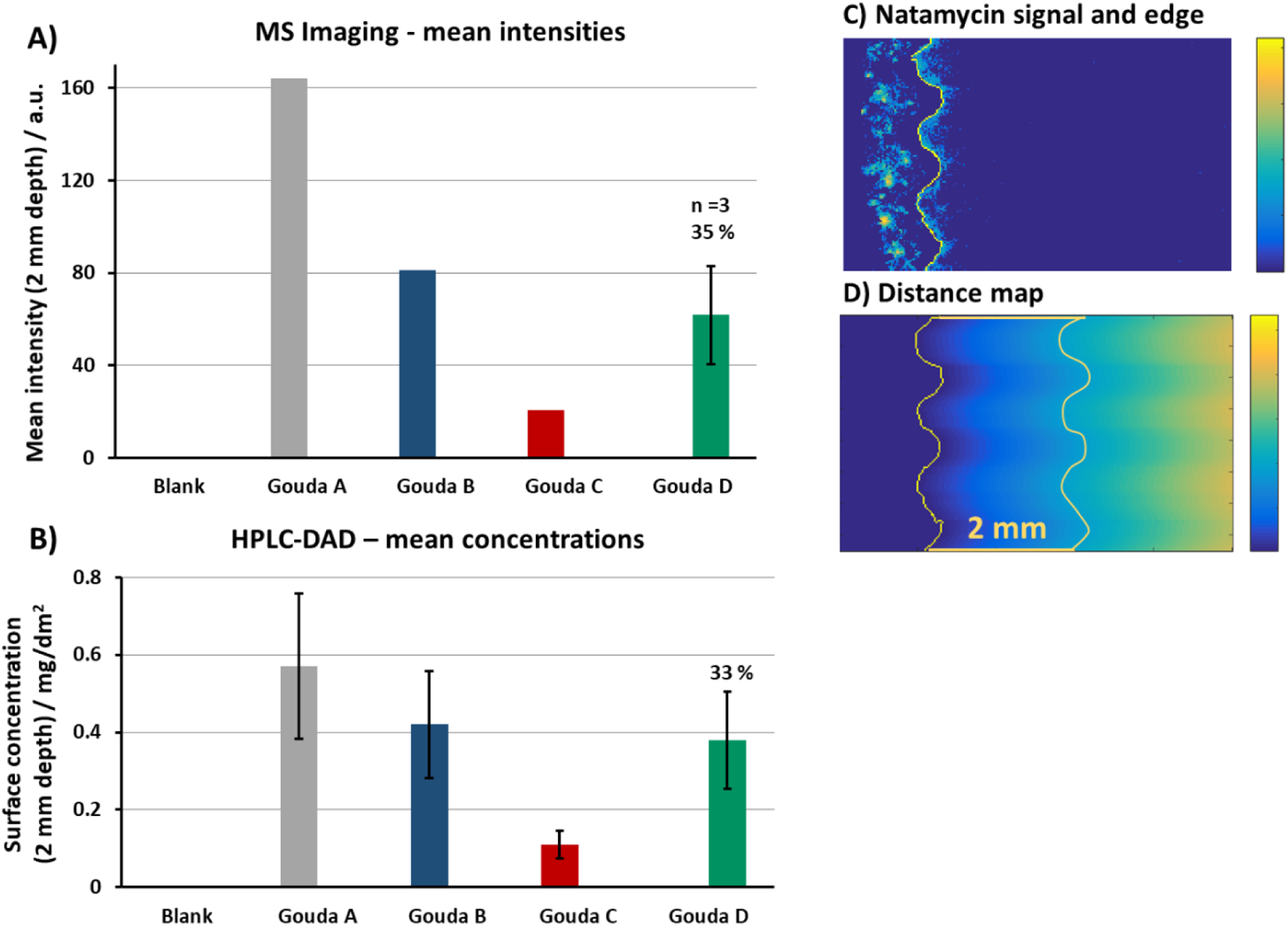
Comparison of natamycin signal intensities among Gouda cheese samples. A: Mean intensities of natamycin of all pixels in the first 2 mm on cheese in the MS imaging experiments for all four cheese samples. Gouda sample D is shown as mean of three serial sections, with a relative standard deviation of 35%. B: Surface concentrations of natamycin in the first 2 mm found by HPLC-DAD measurements after homogenization (reference method). All values have a relative. standard deviation of 33 %. C: Natamycin signal and edge for Gouda D, section 1. D: Distance map with edge and 2 mm line for Gouda D, section 1.

The mean intensities of natamycin from the imaging experiments for the first 2 mm of the four Gouda cheese sections are shown in Figure S12A. For the MALDI –MS imaging results of Gouda D the mean of three neighboring sections is shown with a relative standard deviation of 35%. In Figure S12 C the natamcyin signal for section 1 of Gouda D is depicted with the calculated edge. In Figure S12D the distance map is shown with a second line at 2 mm after the edge. The mean of the natamycin intensities were calculate for all pixels in the first 2 mm after the edge. For comparison, the HPLC-DAD results from the first 2 mm of the cheese, which were measured by LGL are depicted in Figure S12B. The HPLC-DAD results are shown with a relative standard deviation of 33%. This high relative error results from the low intensities and a low recovery rate. The relative standard deviation was calculated with a different line of LC-experiments. The MS imaging results follow the same trend as the homogenate analysis from the HPLC-DAD reference method.

## Footnotes

1 Commission Regulation (EU) 2017/2158 of 20 November 2017 establishing mitigation measures and benchmark levels for the reduction of the presence of acrylamide in food

2 Art. 4 (1) in conjunction with Annex II Cat. 01.7.2 Regulation (EC) No 1333/2008 of the European Parliament and of the Council of 16 December 2008 on food additives

3 § 7 in conjunction with Appx. 1 KäseV (1965)

4 Art. 4 (1) in conjunction with Annex II Cat. 01.7.2 Regulation (EC) No 1333/2008 of the European Parliament and of the Council of 16 December 2008 on food additives

5 § 64(1) Lebensmittel-und Futtermittelgesetzbuch (LFGB) in conjunction with DIN EN ISO 9233-1 and DIN EN ISO 9233-2

6 Art. 11 (1) lit. a Regulation (EC) No 1333/2008 of the European Parliament and of the Council of 16 December 2008 on food additives

